# *Ex vivo* quantification of anti-tumor T-cell activity upon anti-PD-1 treatment in patient-derived lung tumor-on-chip

**DOI:** 10.1101/2023.06.21.545960

**Authors:** Irina Veith, Arianna Mencattini, Martin Nurmik, Isabelle Damei, Christine Lansche, Solenn Brosseau, Giacomo Gropplero, Stéphanie Corgnac, Joanna Filippi, Nicolas Poté, Pierre Mordant, Jimela Tosello, Christine Sedlik, Eliane Piaggio, Nicolas Girard, Jacques Camonis, Hamasseh Shirvani, Fathia Mami-Chouaib, Fatima Mechta-Grigoriou, Stéphanie Descroix, Eugenio Martinelli, Gérard Zalcman, Maria Carla Parrini

**Author notes:** Equal last author contributions.

## Abstract

There is a compelling need for new approaches to predict efficacy of immunotherapy drugs. Tumor-on-chip technology exploits microfluidics to generate 3D cell co-cultures embedded in hydrogels that recapitulate immune and stromal characteristics of a simplified tumor ecosystem. Here, we present the development and validation of lung-tumor-on-chip platforms to quickly and precisely measure *ex vivo* the effects of immune check-point inhibitors on T-cell-mediated cancer cell death, by exploiting the power of live imaging and advanced image analysis algorithms. These tumor-on-chips were generated with patient-derived autologous primary cells isolated from fresh lung cancer samples, opening the path for applications in personalized medicine. Moreover, cancer-associated fibroblasts were shown to impair the response to anti-PD-1, indicating that tumor-on-chips are capable of recapitulating stroma-dependent mechanisms of immunotherapy resistance. This interdisciplinary combination of microfluidic devices, clinically-relevant cell models, and advanced computational methods, can innovatively improve both the fundamental understanding and clinical efficacy of immuno-oncology drugs.

## INTRODUCTION

The lack of adequate model systems is a critical obstacle in the development and deployment of new effective treatments against cancer. It is now well recognized that conventional cell cultures or animal models fail to accurately predict human responses to oncology treatments, as they do not properly mimic human physiopathology, particularly regarding the immune system ^1^. In the last decade, a novel concept has emerged: the use of micro-physiological systems to achieve a rational simplification of the human body. This led to the creation of new research fields, named organ-on-chip (OoC) and tumor-on-chip (ToC) ^2–4^. More specifically, ToC technology exploits micro-fabrication and microfluidics to generate cell co-cultures embedded in 3D hydrogels that mimic the extra-cellular matrix, recapitulating the immune and stromal characteristics of the tumor ecosystem. While ToC field is exponentially growing, current efforts have mainly relied on established, mostly immortalized, cell lines. It is now time to move toward clinically-relevant cell models, such as primary autologous cells, and to prepare the ground for translational applications in personalized medicine.

Clinical oncology is currently undergoing an extraordinary therapeutic revolution driven by the new immunotherapy drugs, in particular immune checkpoint inhibitors (ICI), which can induce impressive long-lasting responses and increase patients’ life expectancy. Melanoma, lung cancer, head and neck, and bladder cancer have most benefited from these immunotherapies, while many clinical trials are still ongoing to show survival improvements in other common cancers. There is a compelling need of new concepts and methods to develop and test immuno-oncology drugs. By definition these new experimental models must be immunocompetent, i.e. they must be able to recapitulate drug effects that rely on the immune components of the tumor microenvironment. In this regard, ToC platforms have great potential that is only waiting to be exploited ^3^. For example, we previously showed that a specific immune response, the antibody-dependent cell-mediated cytotoxicity (ADCC), can be recapitulated in a HER2+ breast ToC treated with trastuzumab (Herceptin) ^5^. Another ToC study revealed a cooperative behavior between cytotoxic T cells in tumor killing ^6^.

In this work we aimed at generating ToC platforms as a novel *ex vivo* experimental paradigm for preclinical studies on ICI responses, in order to pave a new way to address immuno-oncology issues in fully human, controllable and directly observable tumor 3D ecosystems. We chose to focus on lung cancers, the leading cause of cancer-related death worldwide, among which non-small cell lung cancer (NSCLC) is the most frequent lung cancer type (80% of cases). Despite the major improvements achieved by immunotherapy drugs, only 20 to 40% of NSCLC patients benefit from ICI drugs ^7, 8^. What makes non-responder patients resistant to ICI still remains elusive. Undoubtedly, intrinsic cancer cell features, such as PD-L1 expression, tumor mutational burden (TMB) or specific genetic alterations (p53, K-Ras, PTEN, EGFR, STK11, KEAP11 mutations), impact ICI response ^9–11^. However, at the individual level, such biomarkers cannot be reliably used to exclude a patient from immunotherapy, since major responses are still reported in patients with low TMB or low PD-L1 expression ^12, 13^. Besides cancer cells, stromal cell populations, such as cancer-associated fibroblasts ^14^ and endothelial cells ^15^, also contribute to immunotherapy response or resistance. Overall, despite intense research, there is still an unmet medical need for new predictive tools and biomarkers to identify more accurately which patients will derive long-term benefit from ICI. In addition, a crucial challenge is to more accurately understand the mechanisms of primary or secondary resistance to immunotherapy, in order to conceive novel therapeutic interventions capable of switching non-responders to long-term responding patients, by shattering these protection processes.

We reasoned that the study of the cancer-immune interplay, and of its responses to ICI, requires the use of autologous cytotoxic T lymphocytes (CTLs) to avoid any allogeneic reaction. We first used an already established pair of a NSCLC cell line (IGR-Heu) and autologous CTLs (H5B) ^16^, in order to implement robust methods to precisely quantify T-cell-mediated anti-tumor activity upon immunotherapy treatment using ToC platforms. Next, we moved to primary cells, freshly isolated from NSCLC samples, in order to evaluate the possibility to use patient-derived ToCs for personalized immunotherapy response profiling, in a time window of few days, compatible with the decision-making process in clinics. Importantly, our experimental strategy involves the continuous live imaging for two days of the autologous 3D ToC co-cultures, rather than end-point assays, in order to quantify the dynamics of crucial cellular processes within the tumor ecosystem, such as cancer cell apoptosis ^17^ and cancer-immune interactions ^5, 18, 19^. Moreover, the ToC miniaturization allows to use small tumor samples and limited amounts of cells.

This original combination of ToC 3D co-cultures, patient-derived autologous cell models, and advanced computational methods for image analysis, allowed us to develop and validate a novel procedure in order to measure *ex vivo* the effects of immunotherapy treatments on T-cell-mediated anti-tumor activity, opening new avenues for both fundamental and translational research in immuno-oncology.

## RESULTS

### Lung tumor-on-chip (ToC) platforms for personalized immunotherapy response profiling

We conceived a strategy to efficiently generate patient-derived lung ToC, to treat them with immunotherapy drugs and to precisely measure drug response via computational image analysis (Fig. 1A). We used commercially available microfluidic devices (AIM-Biotech), made of plastic and of a gas permeable membrane. The central micro-chamber contains a 3D biomimetic collagen I gel (volume of 3.4 μl), while the two lateral chambers contain medium, to which drugs can be added in static condition or by microfluidic perfusion. Cell populations were embedded in the central collagen gel (2.3 mg/mL): cancer cells (2000 cells/μl density), immune cells (1000 cells/μl density), and cancer-associated fibroblasts (CAFs) (400 cells/μl density). ToC were then imaged under an inverted video-microscope with controlled CO_2_ (5%) and temperature (37°C) for 48 h. The numbers of seeded cells were chosen mainly to offer a proper imaging of the tumor ecosystem and in particular to assess cell/cell interactions; the ToC cell densities were therefore much lower than the *in vivo* real cell densities, but the cell to cell ratios (1:2 immune to cancer, 5:1 cancer to CAF) were reminiscent of physiological situations, taking into account the fact that large variations of CD8+ T-cell proportions have been reported for NSCLC samples (ranging from <2% to 60%) ^20, 21^. For precise quantifications of cell dynamics and interactions, while we initially seeded isolated cells, tumor cells often had the tendency to form multi-cellular clusters, the situation being very heterogeneous depending on each individual patient. Depending on the experiment, live fluorescent dyes were used to selectively pre-stain the different cell populations (CellTrace, red fluorescence) and to monitor apoptotic death (CellEvent Caspase-3/7, green fluorescence). Quantitative image analysis methods were developed to measure parameters that were chosen for their relevance to evaluate immunotherapy responses: death rate of cancer cells, number and time of cancer-immune interactions. The ToC movies were analyzed as pseudo-2D videos since the gel height is relatively small (250 μm). This had the great advantage of avoiding the complexity of 3D tracking algorithms, although the reconstituted lung tumor microenvironments actually have a 3D architecture. Indeed, confocal microscopy confirmed that within the lung ToC the various cell populations were well distributed along the z axis and made many cell-cell physical contacts (Fig. 1B).

**Fig. 1.**
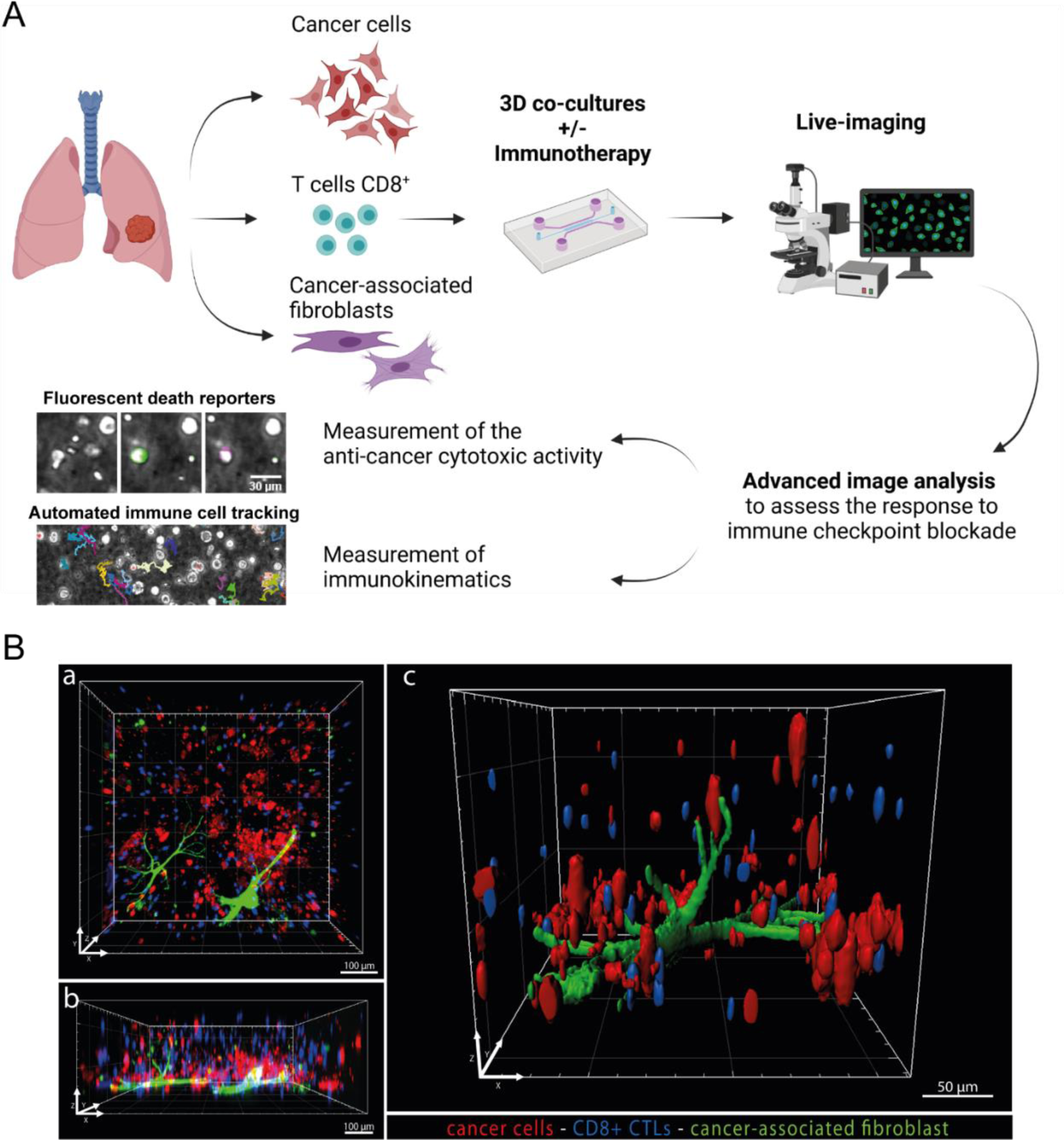
Lung tumor-on-chip (ToC) platforms for personalized immunotherapy response profiling. **A.** Workflow for lung ToC generation and analysis. Cancer cells, T cells and fibroblasts are isolated from the tumor and co-cultured embedded in a biomimetic collagen gel within microfluidic devices. The microfluidic setup allows to perfuse the immunotherapy drugs into the ToC which is live imaged by video-microscopy. Automated advanced methods of image analysis are used to measure the anti-cancer cytotoxic activity and the kinematics of immune cells. **B.** Representative confocal images of the reconstituted 3D lung tumor microenvironment. Autologous cancer cells (IGR-Heu) and CD8^+^ CTLs (H5B) are labeled in red and blue (Cell Trace), respectively. Cancer-associated fibroblasts (CAF07-AD, heterologous) are labeled in green. a, top view. b, lateral view. c, magnified view.

### Autologous lung ToC platforms respond to anti-PD-1 treatment

To develop our model, we first used an already established pair of a NSCLC cell line (IGR-Heu) and its autologous T-cell counterpart (H5B) generated from tumor-infiltrating lymphocytes (TIL) ^16^. The autologous ToC co-cultures of cancer and CTLs were imaged for 48h before and after treatment with an anti-PD-1 immunotherapy drug (nivolumab) (Fig. 2A, Movie 1). Drug injection was achieved by infusing 10 μg/mL anti-PD-1 drug, at 1 μL/min flow rate, starting at 16 h of co-culture. An appropriate microfluidic setup allowed for simultaneous and parallel injection of 3 chips per experiment.

**Fig. 2.**
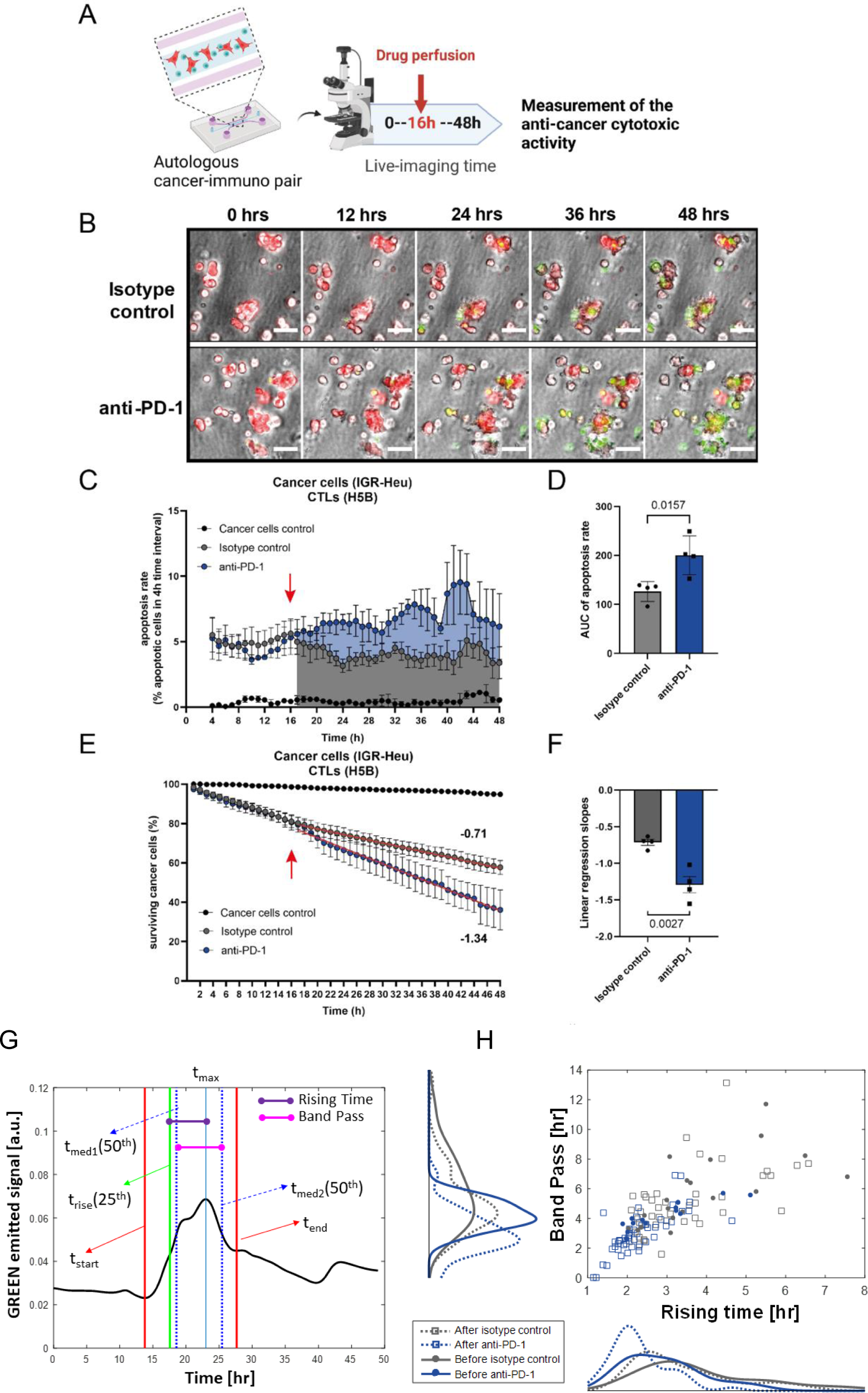
Direct visualization and quantification of *ex vivo* CTL-mediated anti-tumor cytotoxic activity upon anti-PD-1 treatment. **A.** Experimental design. Autologous 3D co-cultures at 2:1 ratio of lung cancer cells (IGR-Heu) and CTLs (H5B) were generated in the central chamber of microfluidic devices and imaged by video-microscopy for 48 h. Anti-PD-1 immunotherapy drug (nivolumab) is perfused in the lateral medium chamber after 16 h of co-culture. **B.** Representative time-lapse images of co-cultures at the indicated time points, before and after the addition of a control isotype antibodies (upper panels) or of anti-PD-1 immunotherapy (lower panels). Cancer cells are stained in red (Cell Trace). CTLs are not stained. A green apoptosis reporter (Cell Event) is added to the medium; cells dying by apoptosis become green. Scale bar, 50 μm. See also supplementary Movie 1. **C.** Quantification of the CTL-mediated anti-tumor cytotoxic activity upon anti-PD-1 treatment. The apoptosis rates of cancer cells, i.e. the percentage of cancer cells dying in 4 h-time-intervals, are computed using the STAMP method ^17^. The averages are computed every 1h using a 4 h-sliding-window. The red arrow indicates the moment of drug injection (16 h). The graph reports means +/- SEM from n=4 independent experiments. **D.** Statistical analysis of apoptosis rates. The areas under the curve from 16h to 48h were measured for the control and treated conditions, from the 4 experiments. Unpaired Student t-test was used. **E.** Quantification of the CTL-mediated anti-tumor cytotoxic activity upon anti-PD-1 treatment. The survival curves of cancer cells, i.e. the percentage of surviving cancer cells calculated with respect to the initial number of living cells, are computed using the STAMP method ^17^. The red arrow indicates the moment of drug injection (16 h). The graph reports means +/- SEM from n=4 independent experiments. **F.** Statistical analysis of survival curves. The linear regression slopes were measured for the control and treated conditions, from the 4 experiments. Unpaired Student t-test was used. **G.** Temporal analysis of death signal at single-cell level. The green signal (Cell Event apoptosis reporter) of one representative dying cell is shown. By automatic signal analysis, characteristic times, t_start_, t_end_, t_max_, t_rise_, t_med1_, and t_med2_ are computed. Then, the Rising Time and the Band Pass are measured. **H.** Plot of the Rising Time and of Band Pass. From 4 conditions: before and after isotype control injection, before and after anti-PD-1 injection. The distribution for each condition, represented by kernel density, are also shown. n=121 cells total. Student t-test showed that differences are statistically significant before versus after anti-PD-1 for Rising Time (p=0.01) and Band pass (p=0.026), but not before versus after isotype control.

Cancer cells were pre-stained in red (Cell Trace). Green apoptosis reporter (Cell Event) was added to the medium at the beginning of co-culture. Red-fluorescent cancer cells dying by apoptosis became green (Fig. 2B, Movie 1). The ToC videos were analyzed by the open-source computational method STAMP (spatiotemporal apoptosis mapper), that we recently developed ^17^. This strategy allowed us to achieve the direct visualization and evaluation of specific antitumor T-cell-mediated cytotoxic activity upon anti-PD-1 treatment. The apoptosis rates of cancer cells (Fig. 2C-D) and the survival curves of cancer cells (Fig. 2E-F), immediately started to diverge upon anti-PD-1 addition, as compared to control isotype antibody control, indicating that CTLs very quickly reacted *ex vivo* to anti-PD-1 blockade.

We analyzed the green fluorescent signal emitted by dying cells at the single-cell level. This green signal is a reporter of intracellular caspase 3/7 activity. Dying cells were manually selected (n=121) and the time profiles of their emitted signals were automatically characterized (Fig. 2G). The distribution analysis showed that after addition of anti-PD-1, but not isotype control, there was a significant decrease in the values of two characteristic signal descriptors, the Rising Time and Band Pass (Fig. 2H) (see Materials and Methods for definitions), indicating that immunotherapy-induced death is characterized by a sharper increase and shorter length of caspase 3/7 activity than ‘natural’ death spontaneously occurring in absence of drug.

Next, we asked the question whether the kinetics of CTL immune cells were modulated by effective anti-PD-1 treatment. For robust tracking of the fast-moving immune cells, phase-contrast ToC videos were generated with high temporal resolution (every 30 seconds for total 6 hours) and with z-stack acquisitions at each position and each time point (Movie 2). Cells were tracked using Cell Hunter software as previously described ^5^.

Cancer-immune interactions were defined as events in which a CTL moved into the circular neighborhood of a cancer cell. This neighborhood was set as 34 μm, twice the sum of the average radius of CTLs (3.4 μm) and cancer cells (13.6 μm) (Fig. 3A-B). The time of interaction between cancer cells and CTLs significantly increased in presence of anti-PD-1, from a median value of 24 min +/- 8 min in the control condition to 29 min +/- 12 min in the presence of immunotherapy treatment (p-value = 0.012) (Fig. 3C). The number of cancer-immune cell interactions during the 6-hour observation time, normalized to the number of cancer cells per video field, significantly increased in presence of anti-PD-1, from average value of 0.07 in the control condition to 0.14 in presence of immunotherapy treatment (p-value <0.0001) (Fig. 3D). In addition, the cell distribution analysis indicated that the addition of anti-PD-1 increased the percentage of both CTL and cancer cells that had high number of interactions (Fig3. E-F): the fraction of CTLs with 3-4 interactions increased from 14% to 19%; the fraction of cancer cells with ≥5 interactions increased from 1% to 7.7%. Moreover, the addition of anti-PD-1 triggered the appearance of a cancer cell fraction (1.5 %) with an extremely high number of immune interactions (>15).

**Fig. 3.**
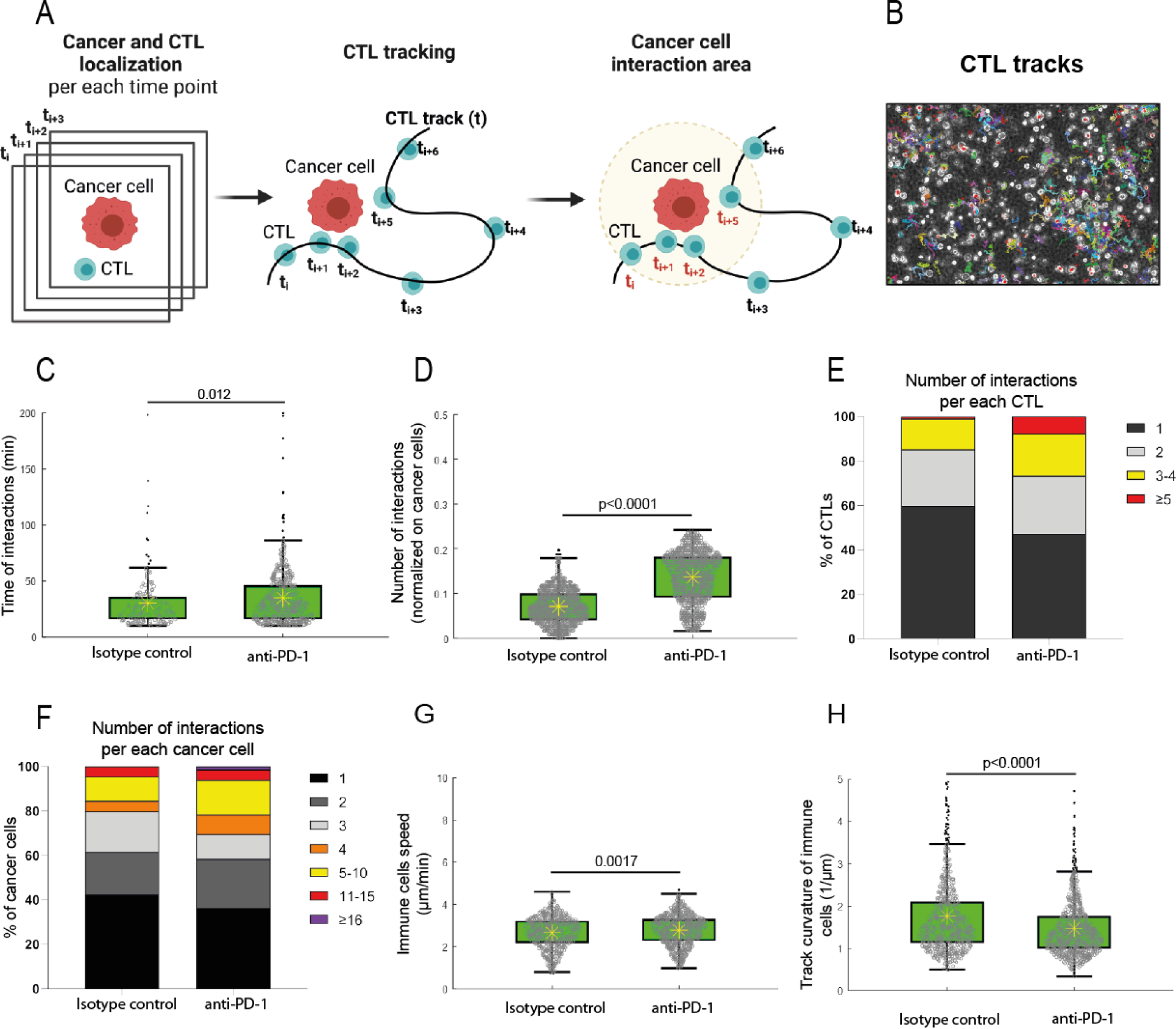
Impact of effective anti-PD-1 treatment on the kinetics of CTL immune cells. **A.** Tracking strategy. ToC videos of lung cancer cells (IGR-Heu) and autologous CTLs (H5B) were acquired with high temporal resolution (every 30 sec) for 6 h. A cancer cell and an immune were considered to interact when their distance was closer than the interaction radius (here defined as 34 μm, twice the sum of the average radius of detected CTLs and cancer cells). See also supplementary Movie 2. **B.** Representative output of Cell Hunter tracking algorithm. **C.** Quantification of the time of interaction between cancer cells and CTLs. n= 669 time events with duration longer than 10 min were counted in total. Statistical significance was assessed using Mann-Whitney test. **D.** Quantification of the number of interactions between cancer cells and CTLs. The number of interactions counted for each cancer cell was normalized by the total number of cancer cell trajectories detected along the video. n=2880 interaction events were counted in total. Mann-Whitney statistical test was used. **E.** Number of interactions per each immune CTL. n=625 interactions in total were counted. **F.** Number of interactions per each cancer cell. n=382 interactions in total were counted. **G.** Quantification of the speeds of immune CTLs. n=1542 speed values were counted in total. Mann-Whitney was used. **H.** Quantification of the track curvatures of immune CTLs. n=1542 curvature values were counted in total. Mann-Whitney was used.

Regarding the immune kinematics, while the average speed of CTLs did not substantially changed in presence of anti-PD-1 (Fig. 3G), the track curvature of CTLs significantly decreased, from a median value of 0.67 μm^-1^ and median absolute deviation of 0.19 μm^-1^ in the control condition to 0.60 μm^-1^ with a median absolute deviation of 0.16 μm^-1^ in presence of immunotherapy treatment (Fig. 3H), indicating that the treatment promotes directionality of CTL movements in 3D setting. Taken together, these results all indicate that autologous lung ToC platforms can reveal drastic and surprisingly rapid cell behavior changes upon anti-PD-1 treatment, as showed by different quantifications: at the cancer cell level (apoptosis rate and apoptotic signal), at the immune cell level (directionality of motility), and at the cancer-immune interaction level (number and duration of interactions). This demonstrates the strong added value of investigating the response to immunotherapy at a tumor ecosystem level and incorporating multi-parametric analysis. The ToC response was fast and precisely quantifiable, supporting the feasibility of immunotherapy response profiling on patient-derived ToC.

### Cancer-associated fibroblasts dampen the response to anti-PD-1 in lung ToC

We previously reported that cancer-associated fibroblasts (CAFs), a major component of tumor stroma, may be involved in immunotherapy resistance in lung cancer patients ^14^. However, the impact of CAFs in immunotherapy response was never experimentally addressed *ex vivo* with patients’ cells. Primary lung CAFs can be easily obtained from tumor samples because CAFs survive relatively well on standard culture dishes. We added allogeneic CAFs (CAF#2, isolated from patient #2) into the autologous cancer-immune co-cultures (IGR-Heu and H5B). Then we treated the cancer-immune-CAF lung ToCs with anti-PD-1 immunotherapy, as described previously (Fig. 4A and 4B, Movie 3). Strikingly, when CAFs were present (5:1 cancer to CAF), the anti-PD-1 treatment was unable to decrease cancer cell survival (Fig. 4C), nor to stimulate apoptotic death (Fig. 4D), indicating that lung CAFs actually promote immunotherapy resistance in the ToC devices.

**Fig. 4.**
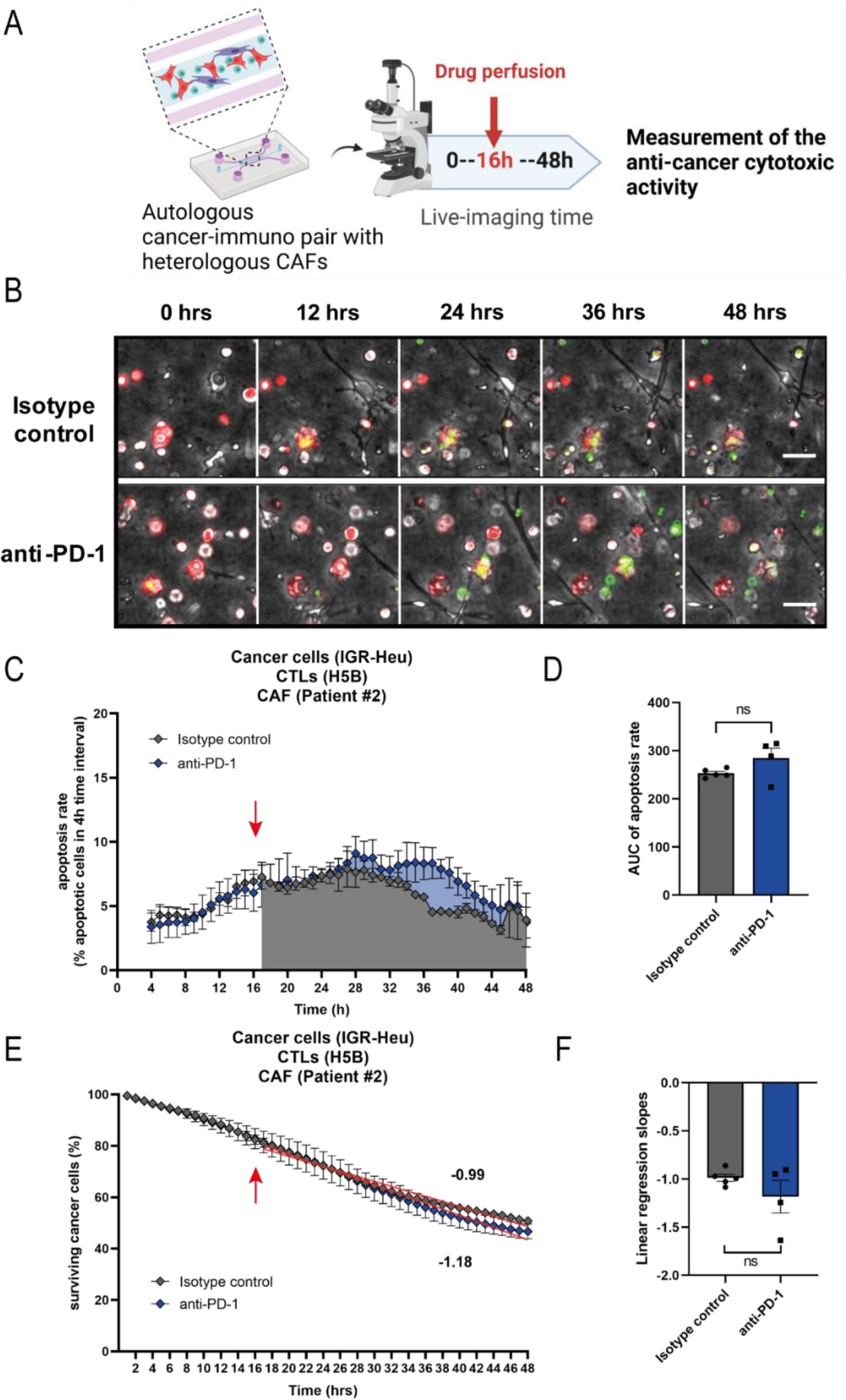
Cancer-associated fibroblasts promote resistance to anti-PD-1 immunotherapy in lung ToC. **A.** Experimental design. Tri-cultures at 2:1:5 ratio of lung cancer cells (IGR-Heu), autologous CTLs (H5B) and heterologous lung CAF (CAF#2) were generated in the central chamber of microfluidic devices and imaged by video-microscopy for 48 h. Anti-PD-1 immunotherapy drug (nivolumab) is perfused in the lateral medium chamber after 16 h of co-culture. **B.** Representative time-lapse images of tri-cultures at the indicated time points, before and after the addition of a control isotype antibodies (upper panels) or of anti-PD-1 immunotherapy (lower panels). Cancer cells are stained in red (Cell Trace). CAFs and CTLs are not stained. A green apoptosis reporter (Cell Event) is added to the medium; cells dying by apoptosis become green. Scale bar, 50 μm. See also supplementary Movie 3. **C.** Quantification of the CTL-mediated anti-tumor cytotoxic activity upon anti-PD-1 treatment. The apoptosis rates of cancer cells, i.e. the percentage of cancer cells dying in 4 h-time-intervals, are computed using the STAMP method ^17^. The averages are computed every 1h using a 4 h-sliding-window. The red arrow indicates the moment of drug injection (16 h). The graph reports means +/- SEM from n=4-5 independent experiments. **D.** Statistical analysis of apoptosis rates. The areas under the curve from 16h to 48h were measured for the control and treated conditions, from the 4-5 experiments. Unpaired Student t-test was used. **E.** Quantification of the CTL-mediated anti-tumor cytotoxic activity upon anti-PD-1 treatment. The survival curves of cancer cells, i.e. the percentage of surviving cancer cells calculated with respect to the initial number of living cells, are computed using the STAMP method ^17^. The red arrow indicates the moment of drug injection (16 h). The graph reports means +/- SEM from n=4-5 independent experiments. **F.** Statistical analysis of survival curves. The linear regression slopes were measured for the control and treated conditions, from the 4-5 experiments. Unpaired Student t-test was used.

These findings support the notion that ToC platforms offer an ideal experimental setting to control the cellular composition of reconstituted tumor ecosystems, and to dissect the contribution of each cell population, such as the CAFs, to immuno-oncology drug resistance.

### Analysis of T-cell plasticity in ToC co-cultures

We reasoned that ToCs might be useful to study T-cell plasticity, in particular their activation and exhaustion status, under controlled co-culture conditions, with or without immunotherapy drugs, and to identify possible alternative targets for ICI treatments. We developed a method to recover cells from ToCs by collagenase digestion, and to analyze T-cell activation and exhaustion markers using multi-color flow cytometry. CD25 and CD69 were selected as activation markers. As exhaustion markers, we looked at PD-1, TIM-3, TIGIT, LAG-3, CD244, CTLA-4 immune checkpoints, and at OX-40, CD137, GITR co-stimulatory receptors (Fig. 5 and Supplementary Fig. S1).

**Fig. 5.**
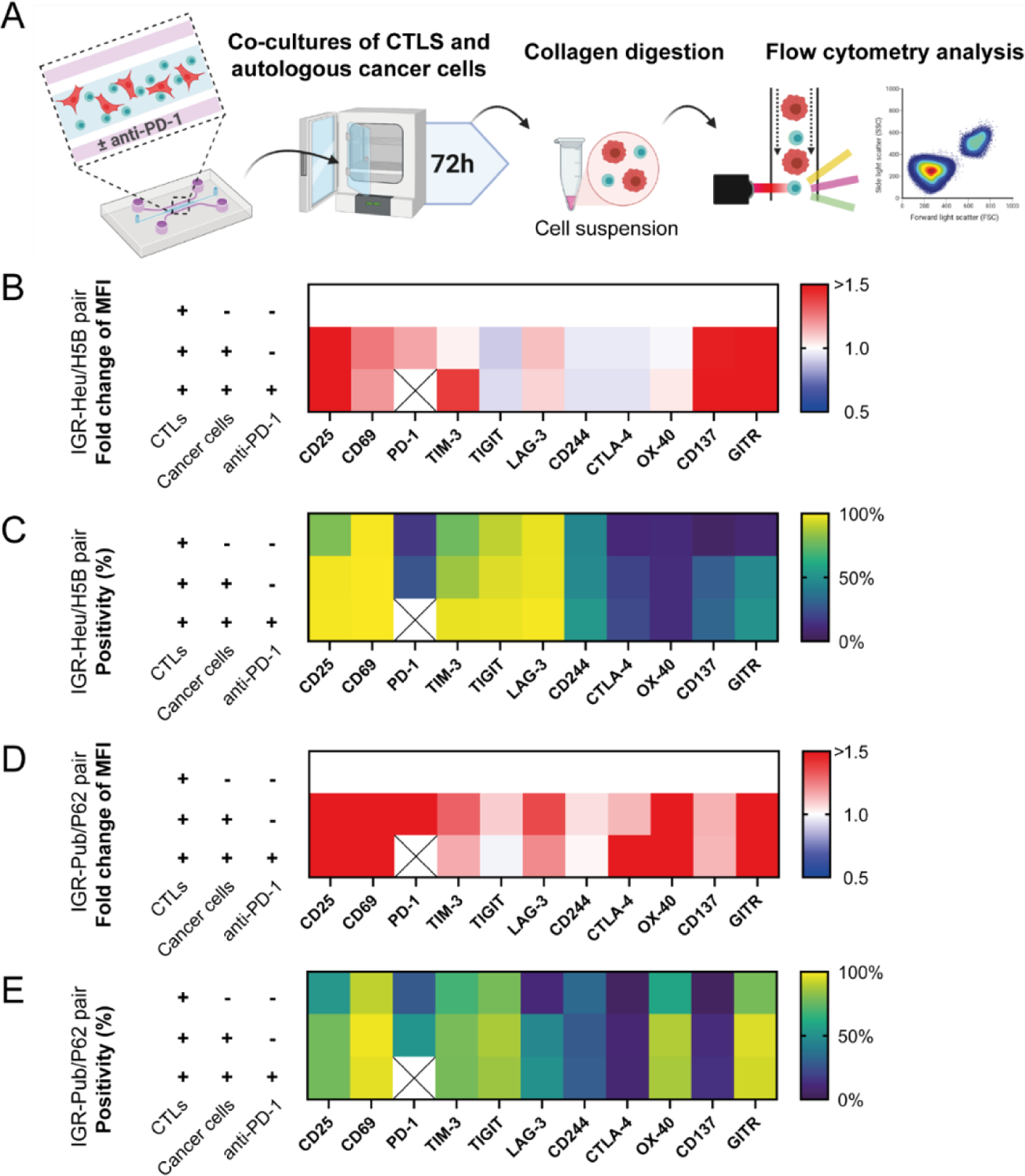
Analysis of T-cell plasticity in ToC co-cultures. **A.** Experimental design. CTLs (H5B or P62) were co-cultured in microfluidic devices with or without autologous cancer cells (IGR-Heu or IGR-Pub), and treated with isotype control or with anti-PD-1 (nivolumab). After 3 days the cells were retrieved from the gel by collagenase digestion, stained and analyzed by flow cytometry. The following markers were measured: CD25, CD69 (activation markers), PD-1, TIM-3, TIGIT, LAG-3, CD244, CTL-4 (inhibitory immune checkpoints), OX-40, CD137, GITR (activatory immune checkpoints). It was not possible to measure PD-1 marker in presence of anti-PD-1 treatment because of antibody competition. Three conditions were assessed: CTLs only, with cancer cells without anti-PD-1, with cancer cells with anti-PD-1. Heat maps report averages from 2 to 4 independent experiments depending on the condition. See Supplementary Fig. S3 for full datasets. **B.** Fold change of specific MFI for CTL markers of H5B cells. The specific MFI for the condition CTLs only is set as 1. **C.** Percentage of positive H5B cells for CTL markers. **D.** Fold change of specific MFI for CTL markers of P62 cells. The specific MFI for the condition CTLs only is set as 1. **E.** Percentage of positive P62 cells for CTL markers.

At basal level, i.e. without co-culture or drug treatment, H5B CTLs displayed a high percentage of positivity (>60%) for many of these markers (CD25, CD69, TIM-3, TIGIT, LAG-3), possibly because these CTLs have been already activated during the *ex vivo* amplification procedure.

After 3 days of co-culture on chip with cancer cells, the H5B CTLs showed a further increase in the expression of activation markers, such as CD25 and CD69, as well as in exhaustion markers, notably PD-1, TIM-3, CD137 and GITR, as assessed by measuring both specific mean fluorescent intensity (MFI) and percentage of positive cells. However, treatment with anti-PD-1 had very mild effects on T-cell expression of both immune checkpoints and co-stimulatory receptors. Only TIM-3 expression appeared to increase, reaching statistical significance for MFI measurements.

To confirm these observations, we tested another established autologous pair: the NSCLC cell line IGR-Pub and the autologous P62 CTL clone ^22^. Again, we observed increased expression in P62 cells of the activation markers CD25 and CD69 upon 3D co-culture with the autologous IGR-Pub cancer cells, as well as of several exhaustion markers, such as PD-1, TIM-3, LAG3, CTL4, OX-40, CD137 and GITR, but no significant modulation by the addition of anti-PD-1 treatment.

These results indicate that when CTLs are already strongly committed into an activation state, by *ex vivo* amplification and/or by co-culture with their target cells, the addition of anti-PD-1 does not substantially impact the expression of immune checkpoints.

### Personalization of ToC using fresh lung cancer samples

In order to achieve personalized immunotherapy response profiling on patient-derived ToC, we optimized a method to freshly isolate primary cells from surgical NSCLC tumor samples by magnetic cell sorting ^23^. Autologous ToC co-culture of cancer and CD8^+^ CTLs were generated the day after patients’ surgery by sequential magnetic-activated cell sorting (MACS) (Fig. 6A) (Movie 4). Cancer and immune cells were directly transferred from *in vivo* environment to *ex vivo* ToC 3D collagen gel, without any culture step on standard dishes. CAFs were amplified in parallel on plastic dishes. We processed 9 lung tumor samples (30-1125 mm^3^ range size), representative of the major NSCLC subtypes: adenocarcinoma, large cell carcinoma, squamous cell carcinoma (Fig. 6B). The cell yields were variable, as well as cell viability (Fig. 6C). Big tumors usually have necrotic cores, leading to low cell viability (see patient #5) and less efficiency in ToC generation. The viability of cancer cells after isolation was the critical parameter to achieve good quality ToCs. We considered ToC generation as a failure when the viability of cancer cells was less than 65% before seeding, since in this condition the co-cultures appeared very heavily influenced by large quantities of initial cell death. In 6 out 8 cases, cell numbers (0.18-1.76 × 10^6^ CD8+ cells, 0.17-4.05 × 10^6^ cancer cells), and survival percentages (68-96%) were perfectly suitable for ToC generation, resulting in an overall 75% success rate.

**Fig. 6.**
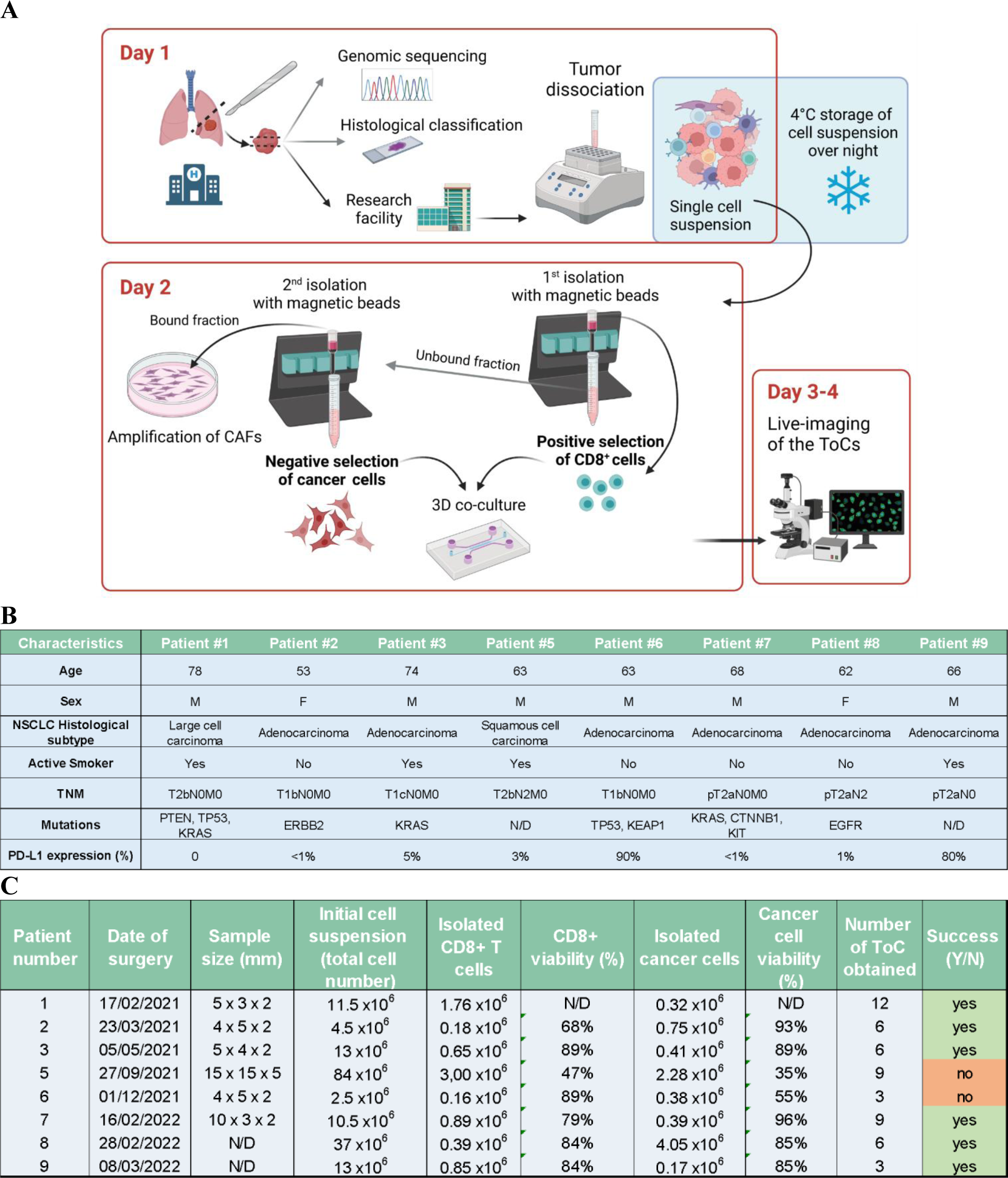
Personalization of ToC using patient-derived cells isolated from fresh lung cancer samples. **A.** Workflow from surgery to tumors-on-chip and live-imaging microscopy. At day 1 the tumor sample is transferred from the hospital to research facility, it is mechanically and enzymatically dissociated; cell suspension is kept over-night at 4°C. At day 2, the three cell types are isolated by magnetic cell sorting: tumor-infiltrating CD8+ CTLs, cancer cells, and CAFs. Autologous 3D co-culture of cancer and CD8+ CTLs are immediately generated within the ToC platforms. Imaging start at day 2 and continue for 48 h during day 3 and day 4. CAFs are amplified in parallel for further use. **B.** Patients’ clinical data. **C.** Efficiency of patient-derived ToC generation. For each patient, it is indicated the tumor sample size, the number of CD8+ and cancer living cells retrieved, the number of ToC obtained, and the overall experiment success.

### Quantification of cancer death induced by autologous CD8+ TILs and by anti-PD-1 immunotherapy in patient-derived lung ToC

Fully automated quantification of cancer death with the original STAMP method was not possible due to the heterogeneity of cancer cell morphology in 3D collagen gels, which is very likely a consequence of the expected patients’ heterogeneity. Moreover, freshly isolated primary lung cancer cells very often constituted multi-cellular aggregates, quickly proliferating, with consequent rapid loss of red Cell Trace dye. To overcome these obstacles, we conceived a TM-STAMP method, in which the segmentation of cancer cell regions was automatically performed in the transmission channel (see Materials and Methods). In order to confirm the reliability of the modified algorithm, we compared original STAMP and TM-STAMP using ToC videos from co-cultures of IGR-Heu cell line and autologous CD8+ H5B T cells. The output results were very similar both in term of time-lapse profiles and of impact of anti-PD-1 drug averaged over the experimental time length (Supplementary Fig. S2), validating the precision of the TM-STAMP method. It is worth to mention that, even though this TM-STAMP version is more adaptable to different ToC morphologies, it is more time-consuming for experimenter and less massive (leading to less data generation).

We analyzed results generated by TM-STAMP from ToC experiments from 6 patients (Fig. 7 and Supplementary Fig. S3). For all patients, the autologous CD8+ TILs had a robust, precisely quantifiable, cytotoxic activity. Interestingly, the cytotoxic activities of the isolated CTLs greatly varied among the patients, reflecting patient heterogeneity (Fig. 7A). For patient #1, we compared the killing effects of different densities of autologous CTLs, at ratio 1:1 and 1:5, highlighting a dose-response effect, as expected (Fig. 7B). ToCs from the other 5 patients (#2, #3, #7, #8, #9) were also treated with anti-PD-1. The addition of anti-PD-1 treatment did not significantly stimulate CTL activity in any patient, but a small trend towards higher average apoptosis rates upon anti-PD-1 treatment was observed in 3 patients (#3, #7, #8). In particular, the time-lapse profiles of apoptosis rates and survival curves of patient 7 suggest that he may be a weak responder. None of these patients ever received anti-PD-1 treatment so far, since they had fully resected early-stage lung cancer with no reimbursed indication yet for adjuvant immunotherapy, so it was not possible to correlate ToC responses with relapse-free survival. In conclusion, these results demonstrate the feasibility to efficiently generate, treat with immunotherapy, and semi-automatically analyze treatment responses in immunocompetent patient-derived ToC, at a time-scale compatible with clinical needs.

**Fig. 7.**
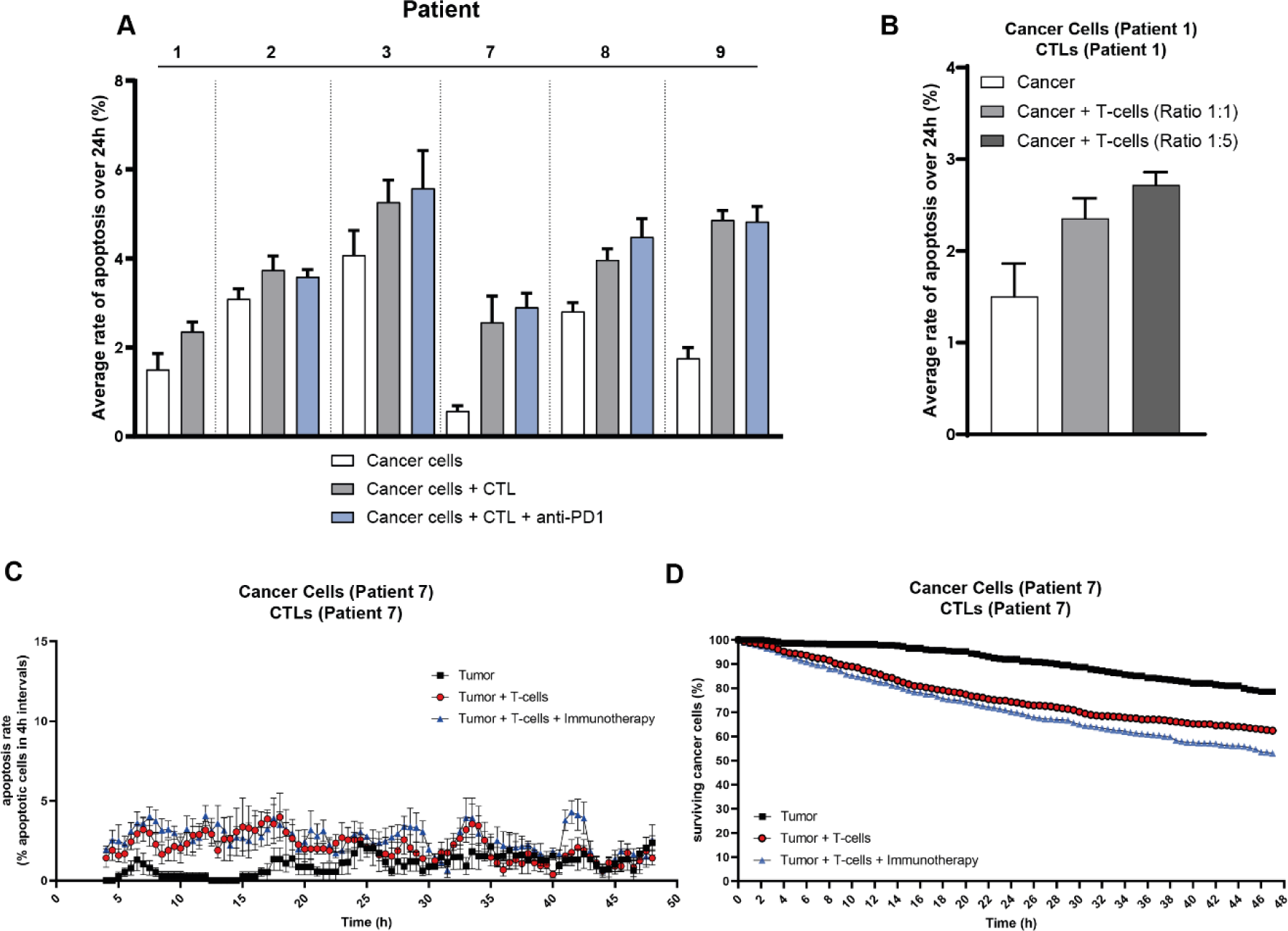
Quantification of CTL-mediated anti-tumor cytotoxic activity in autologous patient-derived lung ToC. **A.** Apoptotic death induced by autologous CD8+ TILs. ToC were generated from 5 NSCLC patients, using primary cancer cells alone or co-cultured with autologous CD8+ TILs at 1:3 ratio. The apoptosis rates of cancer cells were computed over 24 h using the TM-STAMP method and averaged. The graph reports means +/- SEM from 4 view fields for each patient-derived ToC. See also supplementary Movie 4. **B.** Apoptotic death induced by autologous CD8+ TILs as function of CTL density. ToC data shown corresponds to Patient 1 using primary cancer cells alone or co-cultured with autologous CD8+ TILs at 1:1 or 1:5 ratios. The apoptosis rates of cancer cells were computed over 24 h using the TM-STAMP method and averaged. The graph reports means +/- SEM from 4 view fields for ToC derived from patient 1. **C.** Time-lapse representative quantification of CTL-mediated cancer apoptosis rates for Patient 7. The percentage of cancer cells dying in 4 h-time-intervals were computed using the TM-STAMP method. The averages were computed every 1h using a 4 h-sliding-window. The graph reports means +/- SEM from 4 view fields. **D.** Time-lapse representative survival curves for Patient 7. The percentage of surviving cancer cells, calculated with respect to the initial number of living cells, were computed using the TM-STAMP method. The graph reports means +/- SEM from 4 view fields.

## DISCUSSION

Resistance to immunotherapy has dramatic consequences on the outcomes of cancer patients, but currently the only valuable investigation approaches are based on the trial-and-error strategies used in clinical trials and in management of each patient. These empiric approaches are clearly not satisfactory. The success rates in oncology therapeutic clinical trials is extremely low, being recently evaluated to be only at 3.4 % ^24^. Moreover, only 20-40% of lung cancer patients treated by immunotherapy derive long-term benefits, meaning that 80-60% of patients are receiving ineffective treatments, potentially toxic, not to mention their elevated cost leading to the so called ‘financial toxicity’ for Health systems.

In this work, we show that ToC platforms can innovatively improve our understanding of patient-dependent immunotherapy response, as well as resistance mechanisms. Our original ToC procedure for the quantification immunotherapy efficacy has three strengths: it can be personalized, by using patient-derived autologous primary cells directly isolated from fresh tumors; it is precise, because it exploits the power of live imaging and advanced image analysis algorithms; it is relatively fast, in that results can be obtained 3-4 days after patient surgery, a timing compatible with clinical decisions.

Strikingly, we found that the on-chip response to anti-PD-1 injection was already detected 4 hours after drug injection (Fig. 2C and 2E), indicating that reactivation of CTL toxicity develop very fast, which was not known previously. We also showed the feasibility of quantifying CTL toxicity using patient-derived ToCs composed of autologous cancer and immune cells freshly isolated from tumor samples (Fig. 6 and 7). Among our patients, two were clearly non-responders as determined by apoptosis quantification and three showed weak, although noticeable, *ex vivo* response. These observations may be at least partially explained by the *in vivo* intrinsic biological properties of the tumor, since tumors from patients #2 and #7 exhibited CTNNB1 mutations ^25, 26^ or ERBB2 gene amplification ^27^, which have been associated to primary resistance to immunotherapy, while the tumor from patient #3, although being K-Ras mutated, expressed only low amount of PD-L1 (5%). However, we could not validate our predictive power with such a small patient cohort, as none of the patients ever received anti-PD-1 drug, since their early stage tumors were surgically removed, and they did not receive adjuvant post-operative immunotherapy, yet not reimbursed in France in such early stage of the disease.

Other complementary approaches have been proposed for drug screening in personalized therapy design, the main being the organoids and the organotypic cultures (such as tissue slices or tumor fragments). Regarding organoids, despite encouraging experimental evidence supporting the notion that they can recapitulate patient responses in the clinic to chemotherapies or targeted therapies for some cancers ^28–30^, there are doubts on their utility for immunotherapy profiling for lung cancer patients as they require long culture times (weeks) which are incompatible with immune cell survival, and because the overall establishment rate of pure NSCLC organoids is only 17% ^31^. The success rate of our lung ToC was actually much higher: 78 %, although the low number of samples requires caution. Regarding organotypic cultures, a very recent study showed that tumor fragment platforms that preserve the TME architecture, can respond to anti-PD-1 treatment, as assessed by a combination of selected cytokines and T-cell activation markers. Remarkably, in such models, the capacity of tumor-resident T cells to be reactivated *ex vivo* was predictive of clinical response ^32^. Similarly, profiles of secreted cytokines from organotypic tumor spheroids were used to screen for the response of patient tumors to anti-PD-1 therapy ^33, 34^. As compared these organotypic approaches, ToC provide a different type of information since they rely on direct visualization by video-microscopy as potent readout. Indeed, ToC videos produce a wealth of imaging data at single-cell resolution about the dynamics of the TME (e.g., motility, cell-cell interactions, death mitosis, etc.), which can be exploited to deeply investigate the spatio-temporal features of the tumor ecosystem. For example, we recently showed that a deep learning approach is able to correctly identify responding breast ToC cancer-immune co-cultures treated by an immunotherapy drug (trastuzumab), using an atlas of immune cell trajectories ^35^. Moreover, ToCs allow fine TME control of cell parameters (cell types, density, ratio, compartmentalization, etc.), matrix properties (composition, stiffness, porosity, etc.), and the physicochemical environment (dissolved oxygen ^36^, nutrient gradients, etc.) - all of which are very advantageous for addressing the challenges associated with deciphering the resistance mechanisms of immunotherapy.

A weakness of our strategy is that we cannot immediately incorporate the primary CAFs from the same patient in order to generate fully autologous ToCs. In fact, using the magnetic cell sorting protocol, we can isolate three major cell types composing the tumor microenvironment of the same patient: cancer cells, tumor-infiltrating CD8+ CTLs, and CAFs. However, only cancer-immune cell ToC co-cultures can be generated the day after the surgery; so far, the isolation of CAFs requires an amplification step on dish for several days. Amplified primary CAFs can be frozen and used to make heterologous co-cultures. Moreover, it is known that CAFs grown on standard culture dishes give rise to only a specific subset, namely CAF-S1, and cannot account for the huge heterogeneity of CAF identity and functions ^14, 37–39^. More work will be necessary to implement methods capable of isolating the various CAF populations from tumor samples and to incorporate them in ToCs, while also maintaining their original identity.

Despite these limitations, we found that the presence of lung CAFs dampen the response to anti-PD-1 in lung ToCs, experimentally supporting the clinically relevant notion that CAFs play an important role in immunotherapy resistance. Indeed, several *in vivo* studies showed that CAFs, by acting on immune components, can induce an immunosuppressive microenvironment that is likely involved in resistance to immunotherapy and cancer progression. In immunocompetent mouse models, it was reported that FAP+ CAFs are associated with immunosuppression ^40–43^ and induce resistance to common immune checkpoint inhibitors such as anti-PD-1 and anti-CTLA4 through the action of the chemokine CXCL12 ^44^. In breast and ovarian human cancers, CAF-S1 subset displays immunosuppressive functions through a multistep mechanism (*37*, *44*). Notably, in metastatic melanoma and NSCLC, specific CAF-S1 populations, namely the ECM-myCAF, wound-myCAF and TGFβ-myCAF clusters, are enriched at diagnosis in non-responders, indicating that they might be predictive of immunotherapy responses ^14^. In this context, our ToC platforms offer an ideal experimental setting to deeply investigate the CAF-dependent mechanisms of immunotherapy resistance.

In conclusion, this work shows that the use of tumor-on-chip microfluidic platforms is a very valuable asset for the immuno-oncology field. ToC cancer models can be exploited for different purposes: *i)* to investigate the efficacy of new immunotherapy molecules, but also other types of treatments, such as chemotherapy and targeted therapies, also in combination; *ii)* to study the mechanisms underlying resistance to immuno-oncology drugs; *iii)* and to potentially provide in the future new predictive tools to quickly determine the most effective treatment for each patient for a really personalized medicine.

## MATERIALS AND METHODS

### Cell cultures

The IGR-Heu large cell carcinoma cells, the IGR-Pub lung adenocarcinoma cells, and the autologous H5B T-cell line and P62 T-cell clone, respectively, were generated from the same patient in one of our laboratories ^16^. The IGR-Heu and IGR-Pub cancer cells were cultured in DMEM F12 (GIBCO) supplemented with 10% fetal bovine serum (Biosera), 1% of Sodium Pyruvate (Gibco), 1% of Ultroser G (Pall), and 1% Penicillin/Streptomycin (GIBCO). H5B and P62 T cells were cultured in RPMI-1640 (GE Healthcare) supplemented with 10% human AB serum (Institut Jacques Boy, Reims, France), rIL-2 (20 U/ml, Gibco), 1% of Sodium Pyruvate (Gibco) and 0,1% Penicillin/Streptomycin (Gibco). Primary CAFs were cultured in RPMI-1640 (GE Healthcare) supplemented with 10% human AB serum (Institut Jacques Boy, Reims, France), 1% of Sodium Pyruvate (Gibco) and 0,1% Penicillin/Streptomycin (Gibco) and stimulated with irradiated feeder cells as described ^16, 22^. Primary CAFs were cultured in RPMI-1640 (GE Healthcare) supplemented with 10% human AB serum (Institut Jacques Boy, Reims, France), 1% of Sodium Pyruvate (Gibco) and 0,1% Penicillin/Streptomycin (Gibco). T-cell line and clone were periodically tested for tumor cell specificity and to exclude mycoplasma contamination using a qPCR-based method VenorGem Classic, BioValley, #11–1250) or by a plasmid degradation test.

### Isolation of primary cells from fresh lung cancer samples

Fresh tumor samples were provided by the Pathology department of the University Bichat Hospital (AP-HP) and by the Institut Mutualiste Montsouris (IMM) in Paris, France, from patients with NSCLC having undergone standard-of-care surgical resection. Tissue samples were taken from surgical residues available after histopathological analysis and not required for diagnosis. The human experimental procedures follow the Declaration of Helsinki guidelines. Ethical approval was obtained from the institutional review board of the French Society of Respiratory Medicine (Société de Pneumologie de Langue Française, SPLF) (number CEPRO 2020-051) and from the Institutional Review Board and Ethics Committee of Institut Curie Hospital group (CRI-DATA190154). Specimens at IMM were collected under a dedicated protocol approved by the French Ethics and Informatics Commission (EUdract 2017-A03081-52).

Fresh NSCLC samples were stored over night at 4°C in RPMI (10% FBS, 1% Penicillin/Streptomycin). Tumors were cut in small pieces of approximately 2 mm^3^ then incubated with Human tumor dissociation kit (Miltenyi Biotec, #130-095-929) and placed at 37°C in agitation (500 rpm) for 45 min. After enzymatic digestion, the samples were filtered through a 100µm cell strainer (Falcon, Cat#352360) and the remaining pieces smashed with a 1 mm syringe plunger and washed abundantly with Medium A (RPMI 1640, 10% human serum, 10X sodium pyruvate and 0.1% Penicillin/Streptomycin). Necrotic tumors were additionally treated with DNAse I (Sigma, #D5025-15KU, 240 U/mL) for 5min at 37 °C to avoid column clogging during magnetic isolations. After centrifugation the single cell suspension was incubated with RBC lysis buffer (Miltenyi Biotec, #130-094-183) and incubated 2-3 min at room temperature, then washed with Medium A. For CD8+ T cells magnetic isolation we incubated the cell suspension 15 min incubation at 4°C with CD8 microbeads (Miltenyi Biotec, #130-045-201), then cells were passed through LS magnetic columns (Miltenyi Biotec, #130-042-401) installed on a MACS Quadro magnetic stand (Miltenyi Biotec, #130-090-976). To isolate cancer cells, the negative fraction of CD8+ T cells selection was collected and incubated with the Tumor Cell Isolation Kit (Miltenyi Biotec, #130-108-339) for 15 min incubation at 4°C. The cancer cells collected from this second magnetic isolation were directly put in culture in chip with the previously isolated tumor-infiltrated T cells CD8+. In order to retrieve the cancer-associated fibroblasts the cells remaining from the sequential isolations were put in culture in 6-well dishes and amplified.

### Tumor-on-chip preparation

The microfluidic devices were purchased from AIM-Biotech (#DAX-1). Cells were seeded at the final density of 2×10^6^ cells/ml in the central chamber of the DAX-1 chips embedded in a matrix composed of type I rat tail collagen at the final concentration of 2.3 mg/ml (Thermofisher, #A1048301). Each chip can be loaded with 10µl of gel, equivalent to 20000 cancer cells/chip. The targeted cell to cell ratios were 1:2 immune to cancer (effector:target, E:T) and 5:1 cancer to CAF. After addition of the gel in the microfluidic device, in order to allow gel polymerization, chip was incubated in a humidified chamber for 25 min in the incubator (37°C, 5%CO_2_). After gel polymerization, in each lateral chamber 120 µl of T-cell medium were added, supplemented with 6µM CellEvent Caspase-3/7 Green Detection Reagent (Thermofisher, #C10423, green fluorescence). Cancer cell lines (IGR-Heu and IGR-Pub) were pre-stained with 5µM CellTrace Yellow (Thermofisher, #C34567, red fluorescence) in PBS for 25min at 37°C then washed with warm medium and seeded in the gel. Anti-PD-1 antibody (Selleck, #SE-A2002-5MG,) or IgG4 isotype control (Biolegend, #403702) were added at a concentration of 10µg/ml.

### Microfluidic setup

An adapted microfluidic setup was developed in order to inject drugs in the microfluidic devices in real-time while imaging the cells. The microfluidic setup includes, in order: a pressure controller (Fluigent, Flow-EZ), 15ml-tube reservoirs for culture medium, flow control detectors (Fluigent, Flow Unit-M) and the microfluidic device with cells in the video-microscope incubator. The constant flowrate was set at 0.5-2µl/min. All the parts of the microfluidic system are connected through PTFE tubing (Cole Parmer, #06417-11). The tubing was connected to the microfluidic device through custom 3D-printed fluidic adaptors, printed on a DWS 028J+ HR 3D SLA printer with DS3000 clear resin (DWS Systems) and a 100 µm layer height. After printing, the adaptors were rinsed and sonicated in isopropanol for 2 minutes before UV curing for 25 minutes in a UV curing unit (DWS Systems).

### Live cell imaging

The tumor-on-chips were mounted on the motorized stage of an inverted wide-field fluorescence video-microscope (DMi8, Leica), enclosed in an incubation chamber to provide a humid atmosphere with 5% CO2 at 37°C, equipped with a Retiga R6 camera and Lumencor SOLA SE 365 light engine, and piloted by MetaMorph software. 5X objective was used. For each gel, 2-4 positions were acquired. For cancer death analysis, acquisition intervals were 1 frame every 30-60 min for 48 h. Three channels were acquired at each position and each time point: phase-contrast, green and red fluorescence. For green fluorescence, the filters were: excitation 470/40 nm, emission 525/50 nm, dichroic mirror 495 nm. For red fluorescence, the filters were: excitation 560/40 nm, emission 630/75 nm, dichroic mirror 585 nm. For immune cell kinematics, acquisition intervals were 1 frame every 30 sec for 6 h, only in the phase-contrast channel to avoid photo-toxicity. To increase number and length of immune cell tracks, z-stack acquisition mode (6 planes every 5 μm) and the plugin Stack Focuser of ImageJ software were used.

### Flow cytometry

T-cell markers were assessed by flow cytometry after 72 hours of co-culture on-chip with cancer cells, with or without anti-PD-1 treatment. In order to digest the gel and harvest the cells, after a wash with PBS, collagenase I (Millipore, # SCR103, resuspended at 4mg/ml in DMEM F12 and passed through 0.22μm filter) was added to chip lateral chambers for 30 min at 37°C. After collagen digestion, cells were transferred in a 96 well-plate V-bottom (Greiner, Kremsmünster, Austria, #651101) for flow cytometry staining. First cells were stained with Zombie NIR viability staining (Biolegend, #423105, diluted 1:500 in PBS) for 25 min at room temperature. After washing with PBS+ (PBS, 1% FBS, 2mM EDTA) cells were stained with antibody or isotype control mixes for 30 min at room temperature. The antibody mix was composed of anti-CD3 (BD, #561805, diluted 1:60), anti-CD4 (BD, #566804, diluted 1:40), anti-CD8 (BD, #563919, diluted 1:80), anti-CD69 (BD, #747520, diluted 1:40), anti-CD25 (BD, #741365, diluted 1:20), anti-CD279 (BD, #562516, diluted 1:40), anti-TIM-3 (BD, #565567, diluted 1:40), anti-LAG-3 (BD, #565616, diluted 1:40), anti-TIGIT (BD, #747841, diluted 1:40), anti-CTLA-4 (BD, #562742, diluted 1:50), anti-OX-40 (Biolegend, #350012, diluted 1:20), anti-CD137 (BD, #745737, diluted 1:60), anti-GITR (eBiosciences, #15588146, diluted 1:40), anti-CD244 (BD, #550815, diluted 1:20), anti-BTLA (BD, #564800, diluted 1:60). After staining, cells were washed in PBS+ and were analyzed at the ZE5 flow cytometer (Biorad). Compensations were performed using single stained beads (BD biosciences, #552843) for each antibody; for live/dead single staining a mix of live and heat-killed cells (65°C for 5min). Data analysis was performed on FlowJo (v 10.5.2).

### Automatic video analysis

#### Cancer death

The quantifications of cancer apoptosis rates and survival were done using the original STAMP (SpatioTemporal Apoptosis Mapper) method, as previously reported ^17^, for ToC experiments using the cancer cell line (IGR-Heu). For patient-derived ToC experiments, using freshly isolated primary cancer cells, a modification of this method was implemented, named TM-STAMP (TransMission STAMP). In TM-STAMP, the regions occupied by cancer cells, isolated or as aggregates, were automatically segmented at each time frame by using the Circular Hough transform (CHT) ^45^, in the transmission channel instead of red channel. Then, images were merged to create a unique segmentation mask of only the time-lapse regions that were selected in a number of time frames higher than an automatic threshold value estimated through Otsu algorithm ^46^. Since the cancer regions do not move much during the observation time, this unique segmentation mask was applied along the entire video analysis by the STAMP algorithm (Supplementary Fig. S2). One only change was introduced in STAMP code. Since tumor cell aggregates often had heterogeneous shapes, we could not assume cancer cell regions to have circular shapes and we could not anymore define cell background by simply expanding the cell circle in an annulus. Therefore, we applied scaling to the shape manually extracted by the experimenter through morphological operators of dilation ^46^. The resulting background can have diversified shapes according to original shapes of cell aggregates.

#### Single-cell death signal

Video crops of dying cancer cells were manually selected. By applying automatic cell localization by Cell-Hunter software ^5^, the region belonging to the cell was discriminated from the background in order to automatically analyze the average apoptotic green signal (CellEvent Caspase-3/7 reporter) emitted from each cell region. First, a smoothing spline approximation ^47^ was applied over the period 0-48 h. In this way, by smoothing small fluctuations due to errors in cell localization and segmentation, peaks and local minima of green signals were more accurately detected. Then, after detection of maximum intensity value (g_max_) and the corresponding time (t_max_), we applied a local minima finding procedure and retained the two minima, g_start_ and g_end_, whose times are the closest times to t_max_, one before (t_start_) and one after (t_end_) respectively. We computed a baseline value g_0_ as the average values between g_start_ and g_end_. Then, we computed the time, t_rise_, in the range [t_start_,t_max_], as the time at which the green signal reaches the 25^th^ percentile in the range [g_0_,g_max_]. We defined the Rising Time (DT_rise_) the quantity t_max_-t_rise_. We then computed the times t_med1_ and t_med2_ at which the green signal reaches the median value in the range [g_0_,g_max_]. We defined the Band Pass (BP) the quantity t_med2_-t_med1_.

#### Cell kinematics

The quantifications of average speeds and track curvatures of T cells, as well as of the interaction times between cancer and immune cells, were done as previously reported ^19^.

Briefly, the instantaneous speed of T cells along the track was computed as follows:

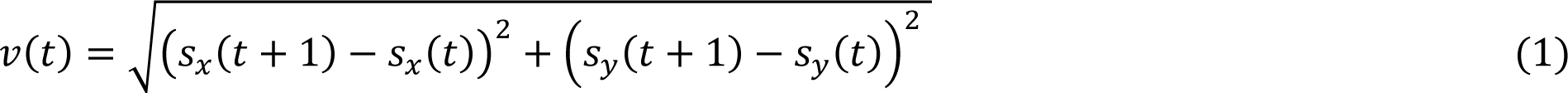

The instantaneous curvature value along the track was computed as follows:

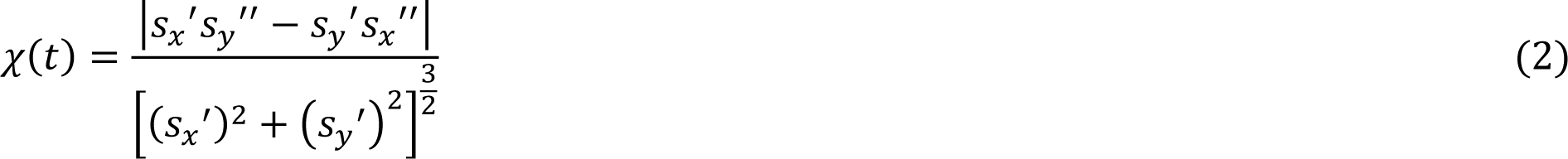

where *s_x_*’ and 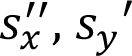 and 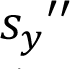, indicate the first and the second-order derivatives along *t* of the *_x_* and *_y_* coordinates, respectively. Subsequently, the average values along the track were computed for each trajectory.

The total number of detected cell tracks was 1600 for CTL and 645 for cancer cells. CTL tracks had an average duration of about 81 min, whereas cancer cells average duration was 137 min. CTL tracks with maximum displacement less than 23 μm and with duration less than 50 min were eliminated and similarly for cancer cell tracks displacement less than 9 μm and duration less than 25 min. Such selection was applied to avoid false cell detection (e.g., phase ring) and to eliminate tracks too short for cell-cell interaction computation.

### Statistical analyses

Statistical analyses were performed using the GraphPad Prism or Matlab software. When the conditions passed the normality Shapiro-Wilk test, a parametric Student t-test was applied. If not, the non-parametric Mann-Whitney or Wilcoxon tests were used. Statistical threshold for significance was set for p-values inferior to 0.05.

## Acknowledgments

We thank the Biological Resources Center (CRB) of the Institut Mutualiste Montsouris (IMM) for providing patient samples, and Jordan Denizeau for technical assistance in tumor sample handling and collection.

## Funding

Fondation ARC pour la Recherche sur le Cancer, PGA1 RF20180206991 and PGA12021010002992_3578.

INSERM, ITMO N°19CR046-00 and ITMO Equipment 2016.

Institut Roche.

CIFRE fellowship to IV founded in part by National Association for Research and Technology (ANRT), on the behalf of the French Ministry of Higher Education and Research, and in part by Institut Roche.

## Author contributions

Conceptualization: IV, GZ, MCP Methodology: GG, SC

Investigation: IV, AM, MN, ID, CL, SB, JF, NP, PM, JT, CS, NG

Visualization: IV, AM, MN, CL

Supervision: EP, JC, HS, FMC, FMG, SD, EM, GZ, MCP

Writing—original draft: IV, AM, MCP

Writing—review & editing: IV, AM, MN, EP, JC, FMC, FMG, SD, EM, GZ, MCP

## Data and materials availability

All data are available in the main text or the supplementary materials.

## REFERENCES

1. Mestas, J. & Hughes, C. C. W. Of mice and not men: differences between mouse and human immunology. J Immunol 172, 2731–2738 (2004).

2. Sung, K. E. & Beebe, D. J. Microfluidic 3D models of cancer. Adv. Drug Deliv. Rev. 79–80, 68–78 (2014).

3. Boussommier-Calleja, A., Li, R., Chen, M. B., Wong, S. C. & Kamm, R. D. Microfluidics: A new tool for modeling cancer-immune interactions. Trends Cancer 2, 6–19 (2016).

4. Sontheimer-Phelps, A., Hassell, B. A. & Ingber, D. E. Modelling cancer in microfluidic human organs-on-chips. Nat. Rev. Cancer 19, 65–81 (2019).

5. Nguyen, M. et al. Dissecting Effects of Anti-cancer Drugs and Cancer-Associated Fibroblasts by On-Chip Reconstitution of Immunocompetent Tumor Microenvironments. Cell Rep 25, 3884–3893.e3 (2018).

6. Ronteix, G. et al. High resolution microfluidic assay and probabilistic modeling reveal cooperation between T cells in tumor killing. Nat Commun 13, 3111 (2022).

7. Paz-Ares, L. et al. Pembrolizumab plus Chemotherapy for Squamous Non-Small-Cell Lung Cancer. N Engl J Med 379, 2040–2051 (2018).

8. Gadgeel, S. et al. Updated Analysis From KEYNOTE-189: Pembrolizumab or Placebo Plus Pemetrexed and Platinum for Previously Untreated Metastatic Nonsquamous Non-Small-Cell Lung Cancer. J Clin Oncol 38, 1505–1517 (2020).

9. Choucair, K. et al. TMB: a promising immune-response biomarker, and potential spearhead in advancing targeted therapy trials. Cancer Gene Ther 27, 841–853 (2020).

10. Cyriac, G. & Gandhi, L. Emerging biomarkers for immune checkpoint inhibition in lung cancer. Semin Cancer Biol 52, 269–277 (2018).

11. Guaitoli, G., Tiseo, M., Di Maio, M., Friboulet, L. & Facchinetti, F. Immune checkpoint inhibitors in oncogene-addicted non-small cell lung cancer: a systematic review and meta-analysis. Transl Lung Cancer Res 10, 2890–2916 (2021).

12. Jardim, D. L., Goodman, A., de Melo Gagliato, D. & Kurzrock, R. The Challenges of Tumor Mutational Burden as an Immunotherapy Biomarker. Cancer Cell 39, 154–173 (2021).

13. Camidge, D. R., Doebele, R. C. & Kerr, K. M. Comparing and contrasting predictive biomarkers for immunotherapy and targeted therapy of NSCLC. Nat Rev Clin Oncol 16, 341–355 (2019).

14. Kieffer, Y. et al. Single-Cell Analysis Reveals Fibroblast Clusters Linked to Immunotherapy Resistance in Cancer. Cancer Discov 10, 1330–1351 (2020).

15. Fukumura, D., Kloepper, J., Amoozgar, Z., Duda, D. G. & Jain, R. K. Enhancing cancer immunotherapy using antiangiogenics: opportunities and challenges. Nat Rev Clin Oncol 15, 325–340 (2018).

16. Echchakir, H. et al. Evidence for in situ expansion of diverse antitumor-specific cytotoxic T lymphocyte clones in a human large cell carcinoma of the lung. Int Immunol 12, 537–546 (2000).

17. Veith, I. et al. Apoptosis mapping in space and time of 3D tumor ecosystems reveals transmissibility of cytotoxic cancer death. PLoS Comput Biol 17, e1008870 (2021).

18. Comes, M. C. et al. The influence of spatial and temporal resolutions on the analysis of cell-cell interaction: a systematic study for time-lapse microscopy applications. Sci Rep 9, 6789 (2019).

19. Mencattini, A. et al. Direct imaging and automatic analysis in tumor-on-chip reveal cooperative antitumoral activity of immune cells and oncolytic vaccinia virus. Biosens Bioelectron 215, 114571 (2022).

20. Lizotte, P. H., et al. Multiparametric profiling of non-small-cell lung cancers reveals distinct immunophenotypes. JCI Insight 1, e89014 (2016).

21. Rakaee, M. et al. Association of Machine Learning-Based Assessment of Tumor-Infiltrating Lymphocytes on Standard Histologic Images With Outcomes of Immunotherapy in Patients With NSCLC. JAMA Oncol 9, 51–60 (2023).

22. Dorothée, G. et al. Tumor-infiltrating CD4+ T lymphocytes express APO2 ligand (APO2L)/TRAIL upon specific stimulation with autologous lung carcinoma cells: role of IFN-alpha on APO2L/TRAIL expression and -mediated cytotoxicity. J. Immunol. 169, 809– 817 (2002).

23. Corgnac, S., Lecluse, Y. & Mami-Chouaib, F. Isolation of tumor-resident CD8+ T cells from human lung tumors. STAR Protoc 2, 100267 (2021).

24. Wong, C. H., Siah, K. W. & Lo, A. W. Estimation of clinical trial success rates and related parameters. Biostatistics 20, 273–286 (2019).

25. Kim, Y. et al. Overexpression of β-Catenin and Cyclin D1 is Associated with Poor Overall Survival in Patients with Stage IA-IIA Squamous Cell Lung Cancer Irrespective of Adjuvant Chemotherapy. J Thorac Oncol 11, 2193–2201 (2016).

26. Pinyol, R., Sia, D. & Llovet, J. M. Immune Exclusion-Wnt/CTNNB1 Class Predicts Resistance to Immunotherapies in HCC. Clin Cancer Res 25, 2021–2023 (2019).

27. Yang, G. et al. First-line immunotherapy or angiogenesis inhibitor plus chemotherapy for HER2-altered NSCLC: a retrospective real-world POLISH study. Ther Adv Med Oncol 14, 17588359221082340 (2022).

28. Vlachogiannis, G. et al. Patient-derived organoids model treatment response of metastatic gastrointestinal cancers. Science 359, 920–926 (2018).

29. Sachs, N. et al. A Living Biobank of Breast Cancer Organoids Captures Disease Heterogeneity. Cell 172, 373–386.e10 (2018).

30. Tiriac, H. et al. Organoid Profiling Identifies Common Responders to Chemotherapy in Pancreatic Cancer. Cancer Discov 8, 1112–1129 (2018).

31. Dijkstra, K. K. et al. Challenges in Establishing Pure Lung Cancer Organoids Limit Their Utility for Personalized Medicine. Cell Rep 31, 107588 (2020).

32. Voabil, P. et al. An ex vivo tumor fragment platform to dissect response to PD-1 blockade in cancer. Nat Med 27, 1250–1261 (2021).

33. Jenkins, R. W. et al. Ex Vivo Profiling of PD-1 Blockade Using Organotypic Tumor Spheroids. Cancer Discov 8, 196–215 (2018).

34. Aref, A. R. et al. 3D microfluidic ex vivo culture of organotypic tumor spheroids to model immune checkpoint blockade. Lab Chip 18, 3129–3143 (2018).

35. Mencattini, A. et al. Discovering the hidden messages within cell trajectories using a deep learning approach for in vitro evaluation of cancer drug treatments. Sci Rep 10, 7653 (2020).

36. Bouquerel, C. et al. Precise and fast control of the dissolved oxygen level for tumor-on-chip. Lab Chip 22, 4443–4455 (2022).

37. Costa, A. et al. Fibroblast Heterogeneity and Immunosuppressive Environment in Human Breast Cancer. Cancer Cell 33, 463–479.e10 (2018).

38. Pelon, F. et al. Cancer-associated fibroblast heterogeneity in axillary lymph nodes drives metastases in breast cancer through complementary mechanisms. Nat Commun 11, 404 (2020).

39. Givel, A.-M. et al. miR200-regulated CXCL12β promotes fibroblast heterogeneity and immunosuppression in ovarian cancers. Nat Commun 9, 1056 (2018).

40. Denton, A. E., Roberts, E. W., Linterman, M. A. & Fearon, D. T. Fibroblastic reticular cells of the lymph node are required for retention of resting but not activated CD8+ T cells. Proc Natl Acad Sci U S A 111, 12139–12144 (2014).

41. Ruhland, M. K. et al. Stromal senescence establishes an immunosuppressive microenvironment that drives tumorigenesis. Nat Commun 7, 11762 (2016).

42. Yang, X. et al. FAP Promotes Immunosuppression by Cancer-Associated Fibroblasts in the Tumor Microenvironment via STAT3-CCL2 Signaling. Cancer Res. 76, 4124–4135 (2016).

43. Zhang, Y. & Ertl, H. C. J. Depletion of FAP+ cells reduces immunosuppressive cells and improves metabolism and functions CD8+T cells within tumors. Oncotarget 7, 23282–23299 (2016).

44. Feig, C. et al. Targeting CXCL12 from FAP-expressing carcinoma-associated fibroblasts synergizes with anti-PD-L1 immunotherapy in pancreatic cancer. Proc. Natl. Acad. Sci. U.S.A. 110, 20212–20217 (2013).

45. Davies, E. R. Machine Vision: Theory, Algorithms, Practicalities. vol. Chapter 10 (Morgan Kauffman Publishers, 2005).

46. Gonzalez, R. & Woods, R. Digital image processing, global edition. Digital Image Processing, Global Edition 19, (2018).

47. Craven, P. & Wahba, G. Smoothing noisy data with spline functions. Numer. Math. 31, 377– 403 (1978).

