## Supplementary Materials for "*Ex vivo* quantification of anti-tumor T-cell activity upon anti-PD-1 treatment in patient-derived lung tumor-on-chip"

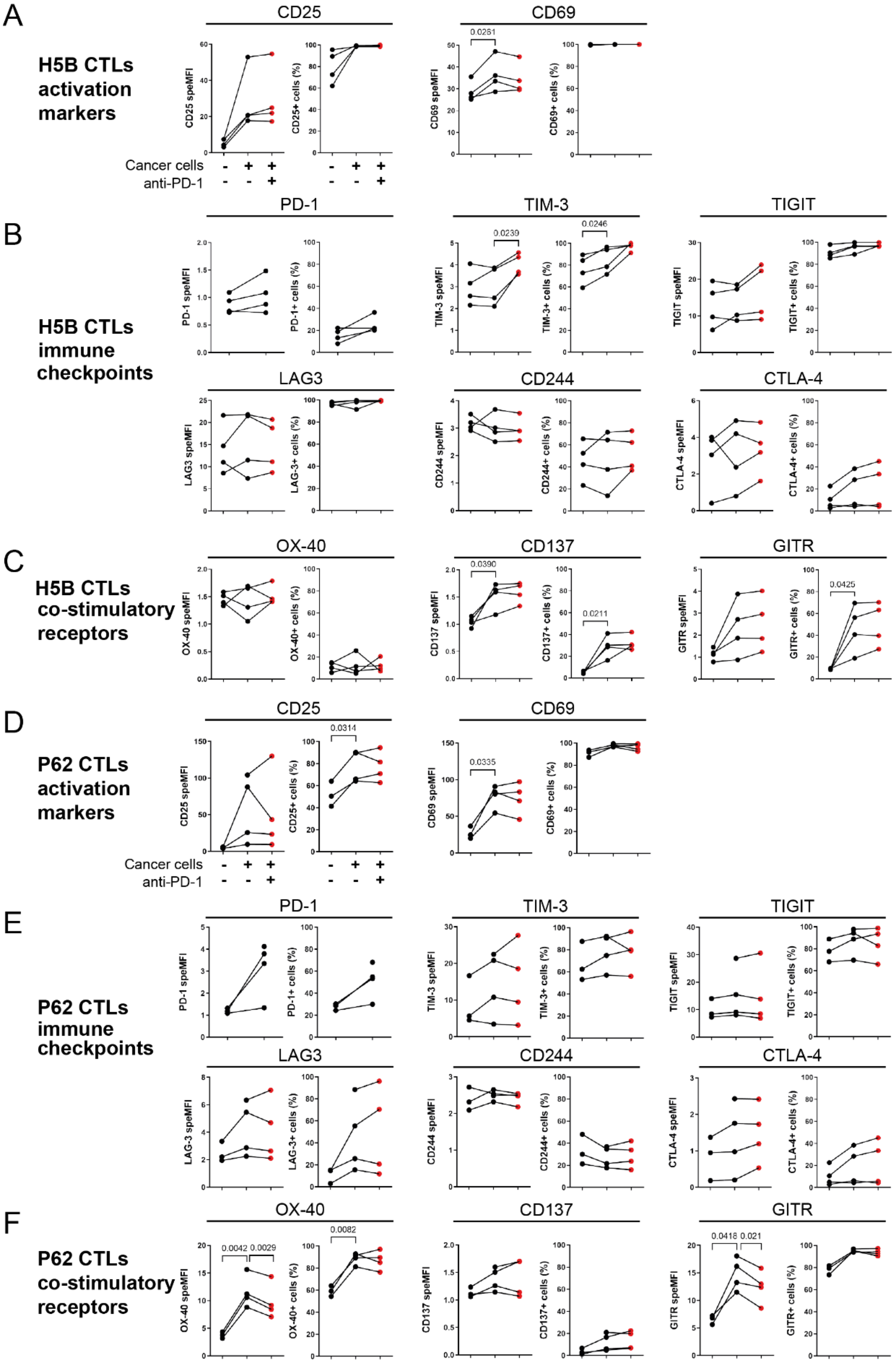

**Supplementary Figure S1. Analysis of T-cell plasticity in ToC co-cultures.**
Full flow cytometry datasets of analysis reported in Fig. 5, from 2 to 4 independent experiments. Red points represent the condition treated with anti-PD-1.
**A.** Specific MFI and percentage of positive cells for activation markers of H5B cells.
**B.** Specific MFI and percentage of positive cells for expression of immune checkpoints on H5B cells.
**C.** Specific MFI and percentage of positive cells for expression of co-stimulatory receptors on H5B cells.
**D.** Specific MFI and percentage of positive cells for activation markers of P62 cells.
**E.** Specific MFI and percentage of positive cells for expression of immune checkpoints on P62 cells.
**F.** Specific MFI and percentage of positive cells for expression of co-stimulatory receptors on P62 cells.
Wilcoxon test was used to determine statistical significance.

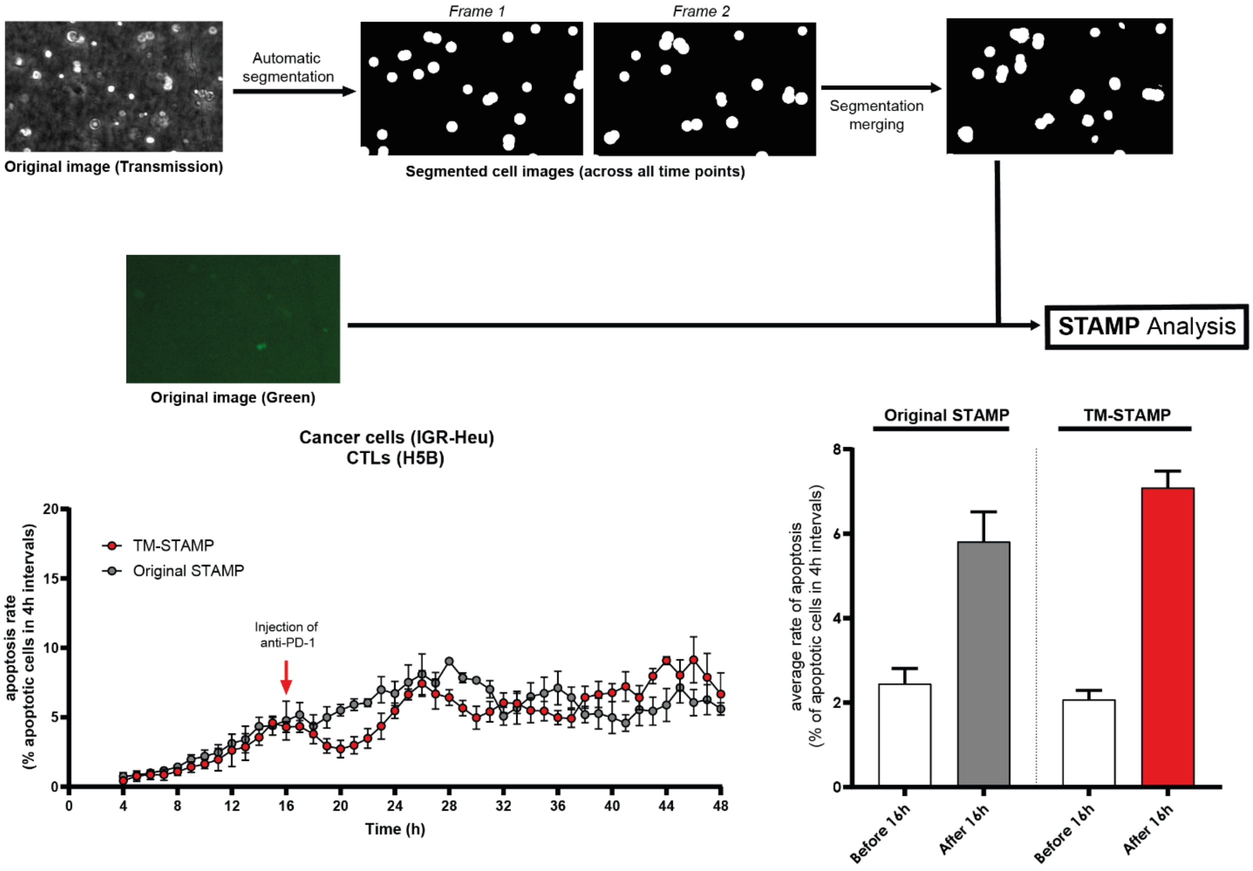


**Supplementary Figure S2. Validation of TM-STAMP method.**

In TM-STAMP areas occupied by tumor cell are automatically identified and segmented based on contrast differences in the transmission channel, instead of red channel used for original STAMP. The segmented cell images are then merged to form the total areas of cellular activity. This unique segmentation mask was applied along the entire video analysis by the STAMP algorithm (see Materials and Methods).

The original STAMP and TM-STAMP methods were compared using videos from one ToC experiment generated using the IGR-Heu/H5B cancer- immune cell pair. The apoptosis rates of cancer cells were computed over 28 h (left graph) and averaged (right graph) over the 16 h before drug injection and over the 16 h after drug injection. The graphs report means +/- SEM from n=4 view fields.

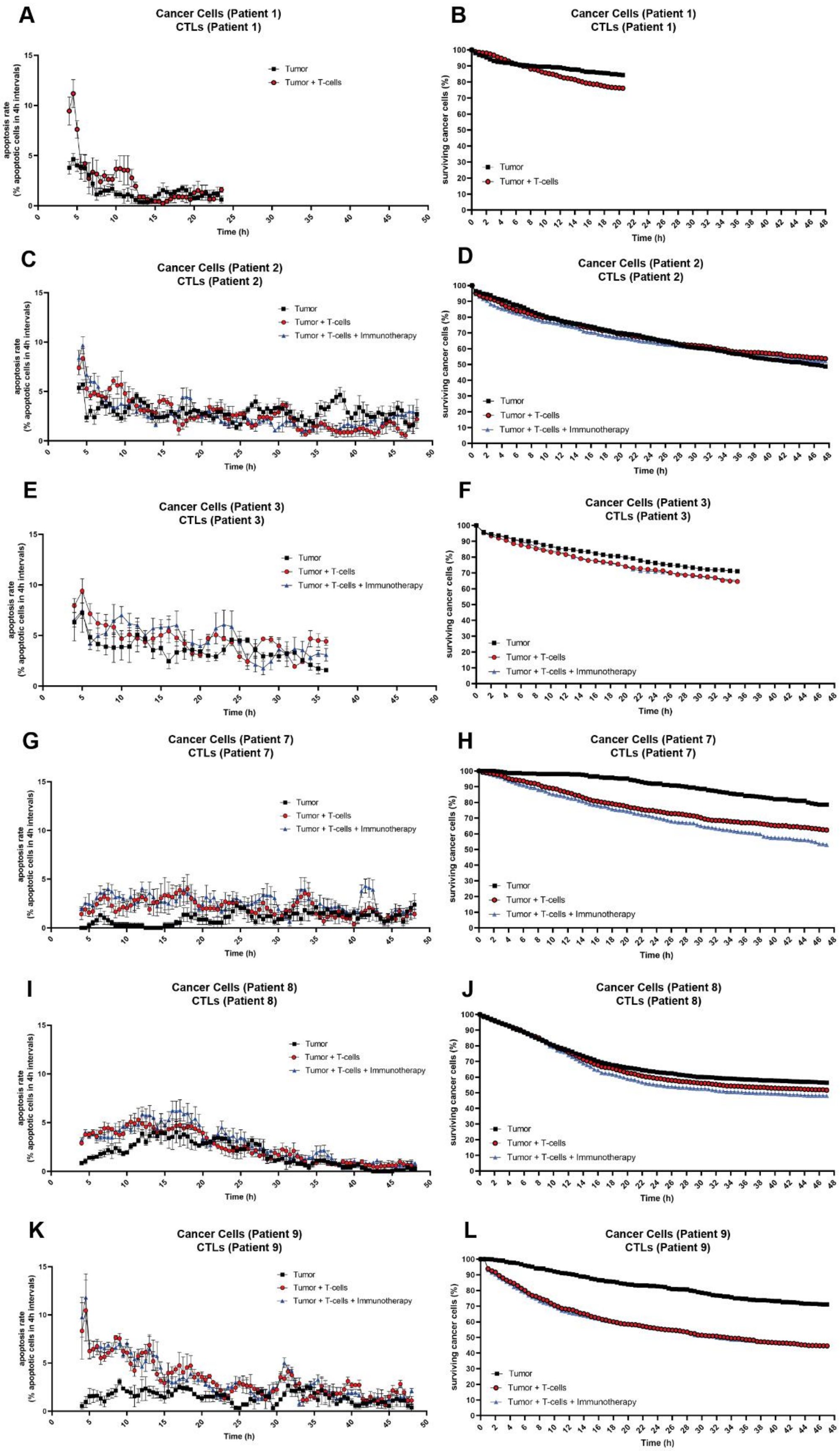
A
Cancer Cells (Patient 1)
CTLs (Patient 1)
Tumor
Tumor + T-cells
apoptosis rate
(% apoptotic cells in 4h intervals)
Time (h)
B
Cancer Cells (Patient 1)
CTLs (Patient 1)
Tumor
Tumor + T-cells
surviving cancer cells (%)
Time (h)
C
Cancer Cells (Patient 2)
CTLs (Patient 2)
Tumor
Tumor + T-cells
Tumor + T-cells + Immunotherapy
apoptosis rate
(% apoptotic cells in 4h intervals)
Time (h)
D
Cancer Cells (Patient 2)
CTLs (Patient 2)
Tumor
Tumor + T-cells
Tumor + T-cells + Immunotherapy
surviving cancer cells (%)
Time (h)
E
Cancer Cells (Patient 3)
CTLs (Patient 3)
Tumor
Tumor + T-cells
Tumor + T-cells + Immunotherapy
apoptosis rate
(% apoptotic cells in 4h intervals)
Time (h)
F
Cancer Cells (Patient 3)
CTLs (Patient 3)
Tumor
Tumor + T-cells
Tumor + T-cells + Immunotherapy
surviving cancer cells (%)
Time (h)
G
Cancer Cells (Patient 7)
CTLs (Patient 7)
Tumor
Tumor + T-cells
Tumor + T-cells + Immunotherapy
apoptosis rate
(% apoptotic cells in 4h intervals)
Time (h)
H
Cancer Cells (Patient 7)
CTLs (Patient 7)
Tumor
Tumor + T-cells
Tumor + T-cells + Immunotherapy
surviving cancer cells (%)
Time (h)
I
Cancer Cells (Patient 8)
CTLs (Patient 8)
Tumor
Tumor + T-cells
Tumor + T-cells + Immunotherapy
apoptosis rate
(% apoptotic cells in 4h intervals)
Time (h)
J
Cancer Cells (Patient 8)
CTLs (Patient 8)
Tumor
Tumor + T-cells
Tumor + T-cells + Immunotherapy
surviving cancer cells (%)
Time (h)
K
Cancer Cells (Patient 9)
CTLs (Patient 9)
Tumor
Tumor + T-cells
Tumor + T-cells + Immunotherapy
apoptosis rate
(% apoptotic cells in 4h intervals)
Time (h)
L
Cancer Cells (Patient 9)
CTLs (Patient 9)
Tumor
Tumor + T-cells
Tumor + T-cells + Immunotherapy
surviving cancer cells (%)
Time (h)

**Supplementary Figure S3. ToC analysis data for all patients.**

**A, B.** Apoptosis rates and survival curves for Patient 1.

**C, D.** Apoptosis rates and survival curves for Patient 2.

**E, F.** Apoptosis rates and survival curves for Patient 3.

**G, H.** Apoptosis rates and survival curves for Patient 7.

**I, J.** Apoptosis rates and survival curves for Patient 8.

**K, L.** Apoptosis rates and survival curves for Patient 9.

The percentage of cancer cells dying in 4 h-time-intervals were computed using the TM-STAMP method. The percentage of surviving cancer cells, calculated with respect to the initial number of living cells, were computed using the TM-STAMP method. The averages were computed every 1h using a 4 h-sliding-window. The observation times varied from 24 h to 48 h, depending on the patient. The graphs report means +/- SEM from 4 view fields.

**Movie S1. ToC co-cultures with lung cancer cells (IGR-Heu) and the autologous CD8$^+$ T-cell line (H5B), treated with anti-PD-1 drug.** Time length, 48 h. Drug is added at 16 h. Cells dying by apoptosis are in green (Cell Event apoptosis reporter). Scale bar, 100 μm.

**Movie S2. High temporal resolution of ToC co-cultures with lung cancer cells (IGR-Heu) and autologous CTLs (H5B), treated with anti-PD-1 drug.** Time length, 6 hours. Drug is added at the start. Dying cells are in magenta (DRQ7 dye). Scale bar, 100 μm.

**Movie S3. ToC tri-cultures with lung cancer cells (IGR-Heu), CTLs (H5B), and CAFs, treated with anti-PD-1 drug.** Time length, 48 h. Drug is added at 16 h. Cells dying by apoptosis are in green (Cell Event apoptosis reporter). Scale bar, 100 μm.

**Movie S4. Patient-derived lung ToC co-culture with cancer cells and CD8+ TILs, treated with anti-PD-1.** Patient #8. Time length, 48 hours. Drug is added at the start. Cells dying by apoptosis are in green (Cell Event apoptosis reporter). Scale bar, 100 μm.
